# Robust Single-cell Matching and Multi-modal Analysis Using Shared and Distinct Features Reveals Orchestrated Immune Responses

**DOI:** 10.1101/2021.12.03.471185

**Authors:** Bokai Zhu, Shuxiao Chen, Yunhao Bai, Han Chen, Nilanjan Mukherjee, Gustavo Vazquez, David R McIlwain, Alexandar Tzankov, Ivan T Lee, Matthias S Matter, Yury Golstev, Zongming Ma, Garry P Nolan, Sizun Jiang

## Abstract

The ability to align individual cellular information from multiple experimental sources, techniques and systems is fundamental for a true systems-level understanding of biological processes. While single-cell transcriptomic studies have transformed our appreciation for the complexities and contributions of diverse cell types to disease, they can be limited in their ability to assess protein-level phenotypic information and beyond. Therefore, matching and integrating single-cell datasets which utilize robust protein measurements across multiple modalities is critical for a deeper understanding of cell states, and signaling pathways particularly within their native tissue context. Current available tools are mainly designed for single-cell transcriptomics matching and integration, and generally rely upon a large number of shared features across datasets for mutual Nearest Neighbor (mNN) matching. This approach is unsuitable when applied to single-cell proteomic datasets, due to the limited number of parameters simultaneously accessed, and lack of shared markers across these experiments. Here, we introduce a novel cell matching algorithm, Matching with pARtIal Overlap (MARIO), that takes into account both shared and distinct features, while consisting of vital filtering steps to avoid sub-optimal matching. MARIO accurately matches and integrates data from different single-cell proteomic and multi-modal methods, including spatial techniques, and has cross-species capabilities. MARIO robustly matched tissue macrophages identified from COVID-19 lung autopsies via CODEX imaging to macrophages recovered from COVID-19 bronchoalveolar lavage fluid via CITE-seq. This cross-platform integrative analysis enabled the identification of unique orchestrated immune responses within the lung of complement-expressing macrophages and their impact on the local tissue microenvironment. MARIO thus provides an analytical framework for unified analysis of single-cell data for a comprehensive understanding of the underlying biological system.

## Introduction

The rapid developments of single-cell technologies have fundamentally transformed our approaches to the investigation of complex biological systems, while potentially influencing clinical decisions. The ability to individually measure the genomic (1), epigenomic (2), transcriptomic (3) and proteomic (4) states at the single-cell level marks an exciting era in biology. Single-cell transcriptomics and targeted-proteomics are the two major approaches commonly used to delineate cell populations and infer functionality or disease states. Single-cell transcriptomics is theoretically able to assess the entire transcriptome of a target cell, with 5-10k unique gene transcripts captured on average for each cell. A key drawback of this method is the relative sparseness of the data generated, particularly for less abundant genes. On the other hand, antibody-based single-cell proteomics has gradually progressed over the years, from the initial detection of a handful of protein targets (5, 6), to about 40 targets via mass cytometry (7), over 100 protein targets via sequencing (8, 9) and most recently, more than 40 protein targets spatially resolved in their native tissue context (10–13). The targeted nature of such approaches requires a careful design, selection, validation and titration of an antibody panel for confident and robust results. Importantly, the features being captured in the biological samples are limited to the antibodies available. Although these factors may limit the number of features that can be measured using targeted single-cell proteomics at any one time, proteomics experiments capture a different spectrum of information than transcriptomics experiments, with following key advantages: first, proteins exert cellular functions, such as signaling cascades, that often define cellular identity, thus allowing a more accurate depiction of the biological state and function, including post-translational events (14, 15); second, although RNA and protein expression can be correlated, RNA counts often do not faithfully represent the final protein machinery expression level in single-cells (16–20); third, due to the limitation of sequencing depth per cell, important but rare transcripts may not be captured in a cell, thus greatly hindering confident cell type annotation (21, 22). In contrast, well-validated antibodies allow robust signal measurements with high dynamic ranges, thus reducing the uncertainties of measurement and chances of false negative or positive events.

Single-cell antibody-based techniques have been widely used, particularly in settings that require robust cell phenotype information or when a specific protein functional readout is necessary. A wide range of single-cell antibody proteomic modalities have now been implemented, including methods like flow cytometry and CyTOF that utilize fluorescent or metal-tagged antibodies to probe large numbers of dissociated suspension cells in a relatively short time (500-10000 cells per second). The parameters assessed include cell surface proteins and intracellular signaling molecules, and samples from different patients or experimental perturbations can be bar-coded and run in the same batch, minimizing variability. Additional methods have recently been developed that allows analysis of proteins in their native spatial contexts (e.g., CODEX, MIBI, IMC), opening a new field of high-parameter tissue biology examination. Sequencing-based approaches such as CITE-seq and REAP-seq can simultaneously probe the RNA and protein levels for each single cell, albeit with the tradeoff of dissociating cells from their original spatial location. Recent methodology developments now allow robust measurements of both nucleic acid and protein information in tissues, although these are currently hindered by either a low number of parameters or poor resolution (23–26).

Given the frequent overlap in proteins measured across dissociated single-cells via sequencing, and intact tissues via antibody-imaging, an orthogonal approach would leverage information from one modality to inform the other. Such an effort would use biological measurements obtained on one modality (e.g. CITE-seq) to inform cells measured using another modality (e.g. CODEX) for a comprehensive assessment of the localization of both proteins and RNAs within tissue samples. Such an approach would be key in inferring either the spatial geolocations of dissociation-based CITE-seq experiments, or the RNA localization of spatial-proteomic CODEX experiments, to enable a better understanding of the complex systems of biological entities.

Several computational approaches for integrative analysis of single-cell data across multiple modalities currently exist (27–30). However, the majority of these methods are tailored toward single-cell sequencing-based analysis, such as scRNA-seq and scATAC-seq, and are not directly compatible with protein-based assays due to differences in the number of parameters and the level of sparsity of the data. The general steps of these methods are the following: Step 1. Project the shared features of the datasets onto a common latent space, from which a cross-dataset distance matrix is constructed; Step 2. Align individual cells greedily via mutual nearest neighbors (mNN); Step 3. Joint embedding of the data and subsequent clustering. Unfortunately, application of this approach to single-cell proteomic datasets can lead to suboptimal results because the number of shared features across proteomic datasets are orders of magnitude smaller than those in single-cell sequencing datasets, and the signals within these limited shared features alone are typically not sufficient to produce high-quality and interpretable pairwise cell matching results. In addition, the intrinsically greedy (and thus at most locally optimal) nature of the mNN matching algorithm limits the ability to fully utilize the correlation structure within the distinct protein features. The first limitation illustrates the necessity of mining the hidden correlations among distinct features, whereas the second roadblock demonstrates the need to optimize the matching objective function to its global optimum. Thus, there is an urgent need for a new strategy specifically designed for matching and integrating single-cell datasets based on limited but robust proteomic parameters.

To meet this need, we have developed *Matching with pARtIal Overlap* (MARIO), a novel algorithm that can robustly match and integrate single-cell datasets based on proteomic measurements. The matching process leverages both shared and distinct features between datasets, and is non-greedy and globally optimized. We additionally developed two quality control steps, the *Matchability Test* and *Joint Regularized Filtering*, to avoid sub-optimal matching and prevent over-integration. Benchmarking of MARIO across various single-cell proteomic data generated from different modalities (CyTOF, CITE-seq and CODEX) and are of cross-species origin (human and non-human primates) demonstrated consistent outperformance of cell-cell matching accuracy over available mNN-based methods. Finally, by applying MARIO, we matched a total of 38,125 macrophages from a CODEX multiplex immunofluorescence lung autopsy dataset to CITE-seq bronchoalveolar lavage fluid (BALF) macrophage cells, and uncovered a spatially orchestrated immune conditioning by complement-expressing macrophages in COVID-19. To make MARIO freely available to the public, we implemented the algorithm in a Python package `MARIO`, along with a R version available online at https://github.com/shuxiaoc/mario-py.

## Results

### Matching and integration of single-cells individually using partially shared features in protein space

There are unique challenges in the implementation of a cell matching algorithm using proteomic information. First, each study is unique and rarely shares identical antibody panels, although a portion of the proteins measured is generally the same. Thus, the matching process must be able to achieve stable pairing of cells with this limited number of features; this is in contrast to transcriptomics data where often several hundred to thousand shared features are available for matching (29, 30). Second, underlying correlations between shared and distinct protein features often exist within and between datasets as a result of panel design and fundamental biological principles. It is therefore pertinent to incorporate information from both shared and distinct protein features. Third, the matching problem should be solved to attain the global optimum rather than a local optimum that is produced by the greedy mNN matching commonly used to align scRNA-seq datasets. Finally, quality control steps are crucial to ensure the accuracy and interpretability of the postulated cell-cell matching results.

To address these challenges, we developed MARIO, a robust framework that accurately matches cells across single-cell proteomic datasets for downstream analysis (Figure 1). MARIO first performs a pairwise cell matching using shared features. To do this, we employ singular value decomposition on shared features to construct a cross-data distance matrix based on the Pearson correlation coefficients of the reduced matrix. An initial cell-cell pairing is then obtained by solving a minimum-weight bipartite matching problem that searches for a distance-minimizing injective map between the two collections of cells. The two datasets are next aligned using this initial matching, and both shared and distinct features of the aligned datasets are projected onto a common subspace using Canonical Correlation Analysis (CCA) (31). This projection is the crux of this methodology as it incorporates the hidden correlations between different proteomic features not shared between the datasets. A cross-dataset distance is then obtained using the canonical scores, and the refined matching is obtained via minimum-weight bipartite matching. By taking the means of the top 10 sample canonical correlations (CCs) as a proxy of matching quality, MARIO then finds the best convex combination weight to interpolate the initial and refined matchings, thus achieving a data-adaptive balancing of the two sources of information.

**Figure 1:**
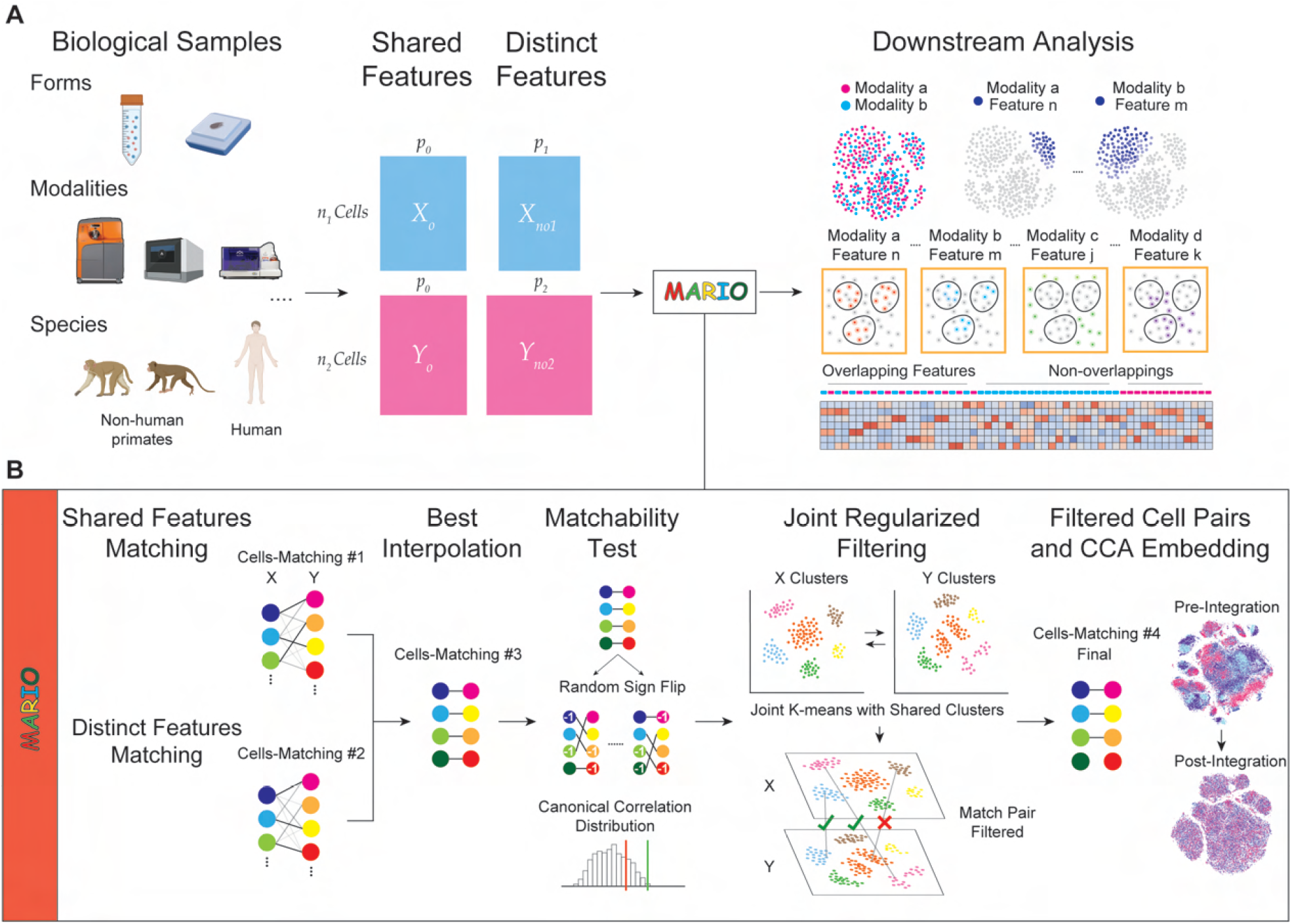
Schematic of the MARIO Analysis Pipeline. **(A)** Single-cell proteomic datasets can be acquired using various modalities, including CyTOF, CITE-seq and CODEX, on different biological samples or species (e.g. human/ non-human primate) with shared underlying biological information. Protein markers are divided into two classes: 1) features captured within both datasets (Shared Features), and 2)markers not shared between the datasets (Distinct Features). Both classes of protein expression matrices serve as inputs to the MARIO algorithm detailed in (B). After the MARIO pipeline, further downstream analysis can be conducted using the combined information integrated across multiple individual experiments. **(B)** A schematic of MARIO algorithm: 1) Individual cells are first subject to matching using the distance matrix constructed using the Shared Features described in (A), before further match refinement using the distance matrix constructed from the Distinct Features such that all features are included. Thereafter, a best interpolation of initial and refined matching will be performed. The dataset then undergoes a matchability test, where random sign flipping is used to validate the statistical rigorosity of MARIO integration using the Canonical Correlation distribution. Subsequently, we perform a cell-cell matching quality control step coined Joint Regularized Filtering, removing spurious cell pairs. Lastly, the matched cells across datasets are jointly embedded into a Canonical Correlation Analysis (CCA) subspace.

After achieving the balanced matching between the two datasets, MARIO next performs a matchability test to determine whether or not the datasets being integrated by the user are suitable for such a joint analysis. It is pertinent that datasets with poor quality or limited underlying correlations are not forcefully paired. The matchability test is performed by flipping the sign of each row of the two datasets with some flipping probability, so that the majorty of underlying correlations (if exists) between the two datasets is abrogated. This process is repeated a number of times to build a distribution of the background CCs of the samples with low underlying correlation. Comparison of the deviation of the sample CCs from the background distribution reveals whether strong underlying information exists to connect the datasets.

Although datasets passing the matchability test are highly correlated, the matching at the individual cell level could still be erroneous if certain rare cell types only exist in one of the dataset or data quality related to specific cell types is inferior. To address these problems, we developed a process termed jointly regularized filtering to automatically filter out low-quality matches without a priori biological knowledge. The filtering process is carried out by optimizing a regularized k-means objective. This objective is a superposition of two parts, where the first part contains individual k-mean clustering objectives for both datasets, and the second part penalizes the Hamming distance between the two individual cluster label vectors and a hypothesized “global” label vector. Use of such a strategy stems from our hypothesis that although the populations being measured in two different experiments may contain modality-specific characteristics (thus the existence of “individual” cluster labels), both originate from a biologically analogous population (thus the existence of a “global” cluster label that is close to the two individual cluster labels). If for a matched pair of cells, the individual labels obtained by joint regularized clustering are not the same, this matched pair is likely spurious and thus disregarded. After this filtering step, the resulting individually matched cells are subject to CCA, and the canonical scores are used as the reduced components for calculating the final embeddings. We implemented generalized Canonical Correlation Analysis (gCCA) to achieve joint embedding of more than two datasets, and subsequently utilized the gCCA sample canonical scores as dimensional-reduced components for calculating and visualizing the final embeddings. Readers are referred to the Materials and Methods section for further descriptions and mathematical details.

### Robust matching and integration of multi-platform and multi-modal single-cell protein measurements with MARIO

We first evaluated the performance of MARIO on two distinctive datasets generated using individual cells isolated from healthy human bone marrow. The first is a sequencing-based CITE-seq dataset consisting of 29,007 cells, stained with an antibody panel of 29 markers (30) and the second is a mass cytometry-based CyTOF dataset consisting of 102,977 cells, stained with an antibody panel of 32 markers (32). Twelve markers (CD11c, CD123, CD14, CD16, CD19, CD3, CD34, CD38, CD4, CD45RA, CD8, and HLA-DR) were common to both datasets. MARIO successfully matched and aligned these two datasets as shown by visual inspection (Figure 2A). The intricate data structures were preserved post-MARIO integration, with clear separation of cells belonging to phenotypically distinctive populations in dimension-reduced t-distributed stochastic neighbor embedding (t-SNE) plots (Figure 2B). The original cell-type annotations based upon the shared low-level annotation (Figure 2B; top left), and on pre-existing annotations from each dataset (Figure 2B; top right and bottom left) were highly conserved after MARIO integration. Subsequent joint clustering of the post-MARIO integrated data using the canonical scores also corroborated in highly accurate cell-type delineation (Figure 2B, bottom right).

**Figure 2:**
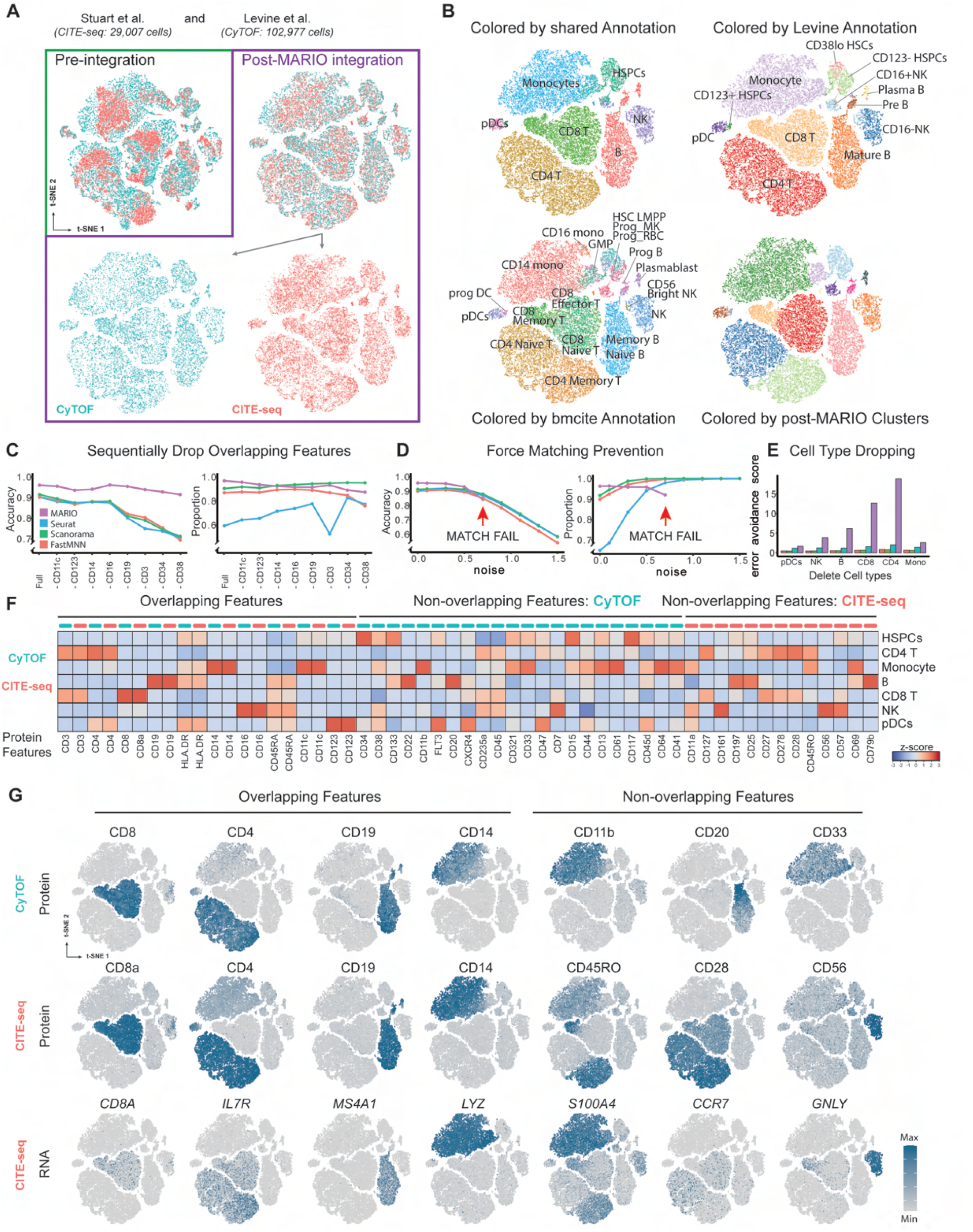
Matching and Integration of CyTOF and CITE-seq Bone Marrow Data using MARIO. **(A)** t-SNE plots of individual cells colored by assay modality, either pre-integration or MARIO integration. **(B)** t-SNE plots of MARIO integrated cells colored by clustering results from (top left to bottom right): High concordance in shared cell types based on annotations from both original datasets; Annotation from Levine et al.; Annotation from Stuart et al.; Clustering result based on CCA scores from MARIO high cell type resolution using information from both assays. **(C-E)** Benchmarking results of MARIO against other mNN-based methods (Purple: MARIO, Blue: Seurat, Green: Scanorama, Red: FastMNN). (C) The matching accuracy (left) and the proportion of cells being matched (right) are tested by sequentially dropping protein features. (D) The matching accuracy (left) and the proportions of cells being matched (right) are measured with increasingly spiked-in noise. (E) The error avoidance score (higher is better) is calculated after dropping each cell type sequentially from the dataset. **(F)** Heatmap of cross modality protein expression levels for the matched cells. **(G)** t-SNE plots of the matched cells with protein/RNA expression levels overlaid based on each of the assays.

We next designed three different scenarios to further characterize the integration performance of MARIO and to compare its performance against the single-cell integration methods Seurat (30), fastMNN (27), and Scanorama (28). In the first case, shared protein markers were removed from each dataset individually (in an accumulative fashion and in alphabetical order) to simulate the distinctive antibody panel designs across potential datasets. MARIO consistently outperformed other methods in terms of matching accuracy, independently of the excluded protein targets (Figure 2C). Thus, MARIO outperformed other methods when used with the plethora of variable experiment-specific antibody panel configurations (full 12-shared panel total accuracy: MARIO, 96.01%; Seurat, 90.29%; fastMNN, 90.22%; Scanorama, 91.46%; dropping 8 shared antibodies: MARIO, 91.45%; Seurat, 70.56%; fastMNN, 69.94%; Scanorama, 71.22%). We additionally evaluated the integration quality among these methods, using metrics including Structure alignment score, Silhouette F1 score, Adjusted Rand Index F1, and Cluster Mixing score, in addition to t-SNE visualizations, based on each method’s post-integration latent space scores (Figure S1A,B).

In the second test, random noise was gradually spiked into the datasets to simulate the variability of intrinsic signal-noise in real world data. The matchability test implemented in MARIO was able to detect and alert the user when data quality was insufficient for confident matching (Figures 2D). In contrast, the elevated noise resulted in an increase in the number of cells being forcefully paired in other tested methods (reaching close to 100%), albeit with low accuracy (ranging from 50% to 80% in accuracy). Given that the other methods are primarily mNN-based and only locally optimized, the higher noise resulted in more erroneous pairs.

In the third scenario, an entire group of cell types was removed from the destination dataset (i.e., the set being matched to) to mimic fluctuations of cell type composition between potential datasets. MARIO outperformed all other tested methods by successfully suppressing the incorrect matching of these missing cell types (Figure 2E; error avoidance scores where larger value indicates better performance for plasmacytoid dendritic cells (pDCs): MARIO, 1.65; mNN methods, 0.42-1.12; natural killer (NK) cells: MARIO, 3.83; mNN methods, 0.40-1.21; B cells: MARIO, 6.18; mNN methods, 0.49-1.15; CD8 T cells: MARIO, 12.67; mNN methods, 0.61-1.57; CD4 T cells: MARIO, 18.89; mNN methods, 0.77-1.99; monocytes: MARIO, 2.60; mNN methods, 0.59-1.39). Given the greedy matching nature of other methods tested, it appears that many of the missing cell types were repeatedly and incorrectly matched with cells from other cell types. This confounding situation is circumvented by the built-in cell-pair filtering function in MARIO.

The precise matching accuracy for CyTOF to CITE-seq cell pairs amongst all the major cell types with MARIO matching was high (Figure S2A): pDCs, 94.57%; NK cells, 98.07%; monocytes, 98.10%; hematopoietic stem and progenitor cells (HSPCs), 76.43%; CD8 T cells, 99.35%; CD4 T cells, 99.64%, and B cells, 98.98%. There was minimal cross-matching, indicative of high accuracy on the single-cell matching level across cell types. Robust matching across two experimental platforms allows the evaluation of differential expression patterns of proteins both shared and unique to these separate experiments. This matching also allows the transcriptome of the single-cells measured using CyTOF to be inferred through the matched CITE-seq pairs. We confirmed that the expression patterns of cell type-specific markers were in good agreement between CyTOF proteins, CITE-seq proteins, and CITE-seq RNA transcripts (Figure 2F, G and Figure S2B, C). Moreover, the expression pattern of CD45RO protein and *S100A4* and *CCR7* RNAs from CITE-seq assisted the delineation of memory and naive CD4 T cell subtypes in the integrated dataset, which was individually unavailable for manual annotation in the CyTOF dataset alone. Therefore, this integrated analysis better defines cell states than do these modalities individually.

We subsequently evaluated the performance of MARIO on two healthy human peripheral blood mononuclear cell (PBMC) datasets measured by CITE-seq and CyTOF. Fifteen proteins (CD11b, CD127, CD14, CD16, CD19, CD25, CD27, CD3, CD4, CD45RA, CD45RO, CD56, CD8a, HLA-DR and PD-1) were common across these two datasets. MARIO successfully integrated the two datasets (Figure S3A) and resulted in accurate cell type matching (Figure S3B; NK cells, 89.93%; naive CD4 T cells, 94.33%; memory CD4 T cells, 90.25%; dendritic cells (DCs), 79.66%; CD8 T cells, 98.69%; monocytes, 96.46%; and B cells, 97.94%). Our results reveal that the expression of key genes on both protein (CyTOF and CITE-seq) and RNA (CITE-seq) levels are in high agreement with their corresponding phenotypic cell-of-origin assignments (Figure S3C). Further benchmarking using the three cases described above showed similar superior matching accuracy for MARIO regardless of antibody panel setup (Figure S4A; full 15-antibody shared panel total accuracy: MARIO, 90.62%; Seurat, 87.55%; fastMNN, 87.27%; Scanorama, 87.39%; dropping 8 shared antibodies total accuracy: MARIO, 86.34%; Seurat, 80.10%; fastMNN, 80.04%; Scanorama, 81.03%). In evaluation of suppression of over-integration due to poor quality data, mNN methods force matched almost all cells with accuracy below 70%, whereas MARIO alerted the user of poor data quality (Figure S4B). Thirdly, integration with MARIO, but not with mNN methods, was robust even with extensive cell type composition changes (Figure S4C; error avoidance scores for monocytes: MARIO, 1.94; mNN methods, 0.53-1.37; B cells: MARIO, 4.53; mNN methods, 0.56-1.37; DCs: MARIO, 1.13; mNN methods, 0.31-0.93; NK cells: MARIO, 2.54; mNN methods, 0.43-1.17; CD8 T cells: MARIO, 4.83; mNN methods, 0.46-1.01; memory CD4 T cells: MARIO, 3.97; mNN methods, 0.38-0.85).

### Cross-species integrative analysis reveals species and stimuli-specific immunological responses

Non-human primates (NHP) are a cornerstone of biomedical research, enabling the rapid investigation of diseases and host responses in a system highly analogous to humans as demonstrated for rapid disease modeling and vaccine development during the recent COVID-19 pandemic (33). Nonetheless, animal models do not fully recapitulate all host responses in humans (34, 35). Given the increasing amount of singlecell proteomic studies in NHP models of disease (36–39), the ability to identify common and different responses to diseases is essential to appreciate host immune response at scale. Given the major commonalities of host immune compositions across NHPs and humans, we postulated that MARIO would be able to effectively integrate human and NHP datasets to reveal underlying common immune coordination and differential responses.

We performed MARIO matching of four CyTOF datasets from studies in which 1) human whole blood cells were isolated from individuals challenged with H1N1 virus (40), consisting of 102,147 cells, 2) human whole blood cells were stimulated with IFN*γ* (37), consisting of 114,175 cells, 3) rhesus macaque whole blood cells were stimulated with IFN*γ*, consisting of 112,218 cells, and 4) cynomolgus monkey whole blood cells were stimulated with IFN*γ*, consisting of 91,409 cells (Figure 3A, B). Dataset 1 was generated using 42 markers, and datasets 2, 3, and 4 were generated using 39 markers. We observed a high degree of concordance between cell types when visualizing the human-human and human-NHP datasets via t-SNE using MARIO integrated canonical scores (Figures 3A, B). MARIO cell-type assignment accuracies were high (Figure 3C). For dataset 1 to dataset 2, accuracies were as follows: B cells, 96.96%; CD4 T cells, 98.80%; CD8 T cells, 98.22%; monocytes, 99.66%; neutrophils, 99.51%; NK cells, 98.39%. For dataset 1 to dataset 3, accuracies were as follows: B cells, 86.76%; CD4 T cells, 97.22%; CD8 T cells, 91.75%; monocytes, 97.85%, neutrophils, 97.99%; NK cells, 86.42%. For dataset 1 to dataset 4, accuracies were as follows: 1 to 4: B cells, 91.90%; CD4 T cells, 96.49%; CD8 T cells, 92.53%; monocytes, 95.14%, neutrophils, 96.10%; NK cells, 80.78%. There were minimal differences, as measured using Euclidean distance, between paired cells calculated by canonical scores (Figure 3D).

**Figure 3:**
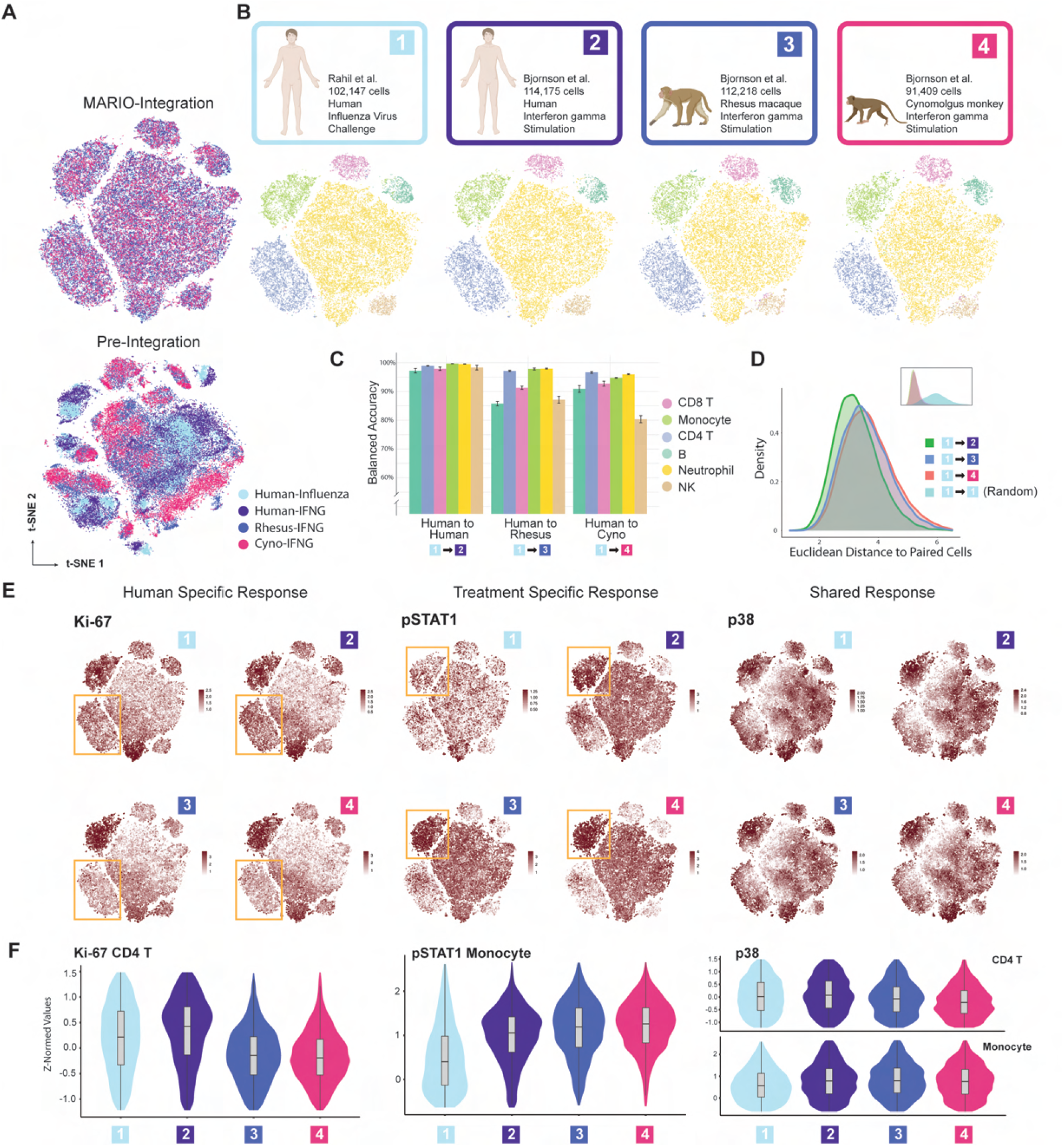
MARIO enables Cross-species and Stimuli Integrative Analysis. **(A)** t-SNE plots of the four datasets, pre- and post-MARIO integration, colored their origin. **(B)** t-SNE of MARIO integrated plots from each individual dataset, colored by cell type. **(C)** Balanced accuracy for each cell type after MARIO matching, for cells from Rahil et al. to other datasets. **(D)** Euclidean distance of canonical correlations for pairs of matched versus random cells between Rahil et al. to other datasets. **(E)** t-SNE plots with expression levels of Ki-67, pSTAT1 and p38 across the four datasets. **(F)** Violin plot of the normalized expression levels of Ki-67, pSTAT1 and p38 across the four datasets for the specified cell types: CD4 T cells and monocytes.

Successful application of MARIO for robust matching and integration across three species and two stimulation conditions allowed us to investigate intrinsic differences in cell type-specific immune responses across humans and NHPs. We observed an increase in proliferation of CD4 T cells in human blood cells after both influenza viral challenge and IFNγ stimulation, as marked by the upregulation of Ki-67, but no increase proliferation was detected after stimulation of NHP blood cells (Figure 3E and F). We also observed the upregulation of pSTAT1, particularly in monocytes, in human and NHP samples treated with IFN*γ* but not after influenza challenge (Figure 3E and F). These results are consistent with previous observations (41–43). Finally, there was an increased p38 expression in all cell types across all samples, reflective of the conserved functionality of p38 during cell inflammatory and stress responses (44, 45). Our benchmarking results showed superior matching accuracy using MARIO regardless of antibody panel setup. When using 39 shared antibodies, the total accuracy was 93.26% for MARIO, 86.20% for Seurat, 84.89% for fastMNN, and 85.83% for Scanorama; when eight shared antibodies were dropped, the total accuracy for IFN*γ* treatment was 86.79% for MARIO, 80.88% for Seurat, 77.89% for fastMNN, and 82.23% for Scanorama (Figure S5A). In the analyses with spiked-in noise, mNN methods forced matching almost 100% of cells with accuracy below 70% with increased noise added, whereas MARIO alerted the user of insufficient information for matching (Figure S5B). MARIO, unlike the mNN methods we tested, was robust in resisting cell-type composition changes (Figure S5C; error avoidance scores, B cells: MARIO, 1.36; mNN methods, 0.51-1.07; NK cells: MARIO, 2.75; mNN methods, 0.52-1.01; neutrophils: MARIO, 2.01; mNN methods, 0.41-1.02; CD8 T cells: MARIO, 1.52; mNN methods, 0.63-0.96; CD4 T cells: MARIO, 1.47; mNN methods, 0.43-0.93; monocytes: MARIO, 1.64; mNN methods, 0.52-1.19)

We similarly applied this strategy to data from IL-4-stimulated human and NHP whole blood cells, and compared them to human influenza viral challenge blood cells (Figure S6A, B). Upon IL-4 stimulation, we saw an upregulation of Ki-67 in human CD4 T cells but not NHP cells, much akin to IFN*γ* stimulation (Figure S6C), and high expression of pSTAT1 in monocytes of IL-4-stimulated blood cells but not in human blood cells challenged with influenza (Figure S6C). In line with IFN*γ* stimulation, the p38 response was consistent across species and treatments. Our results consistently showed superior matching accuracy using MARIO regardless of antibody panel setup. When using 39 shared antibodies, the total accuracy was 89.60% for MARIO, 87.75% for Seurat, 88.30% for fastMNN, and 86.76% for Scanorama; when eight shared antibodies were dropped, the total accuracy was 87.16% for MARIO, 82.72% for Seurat, 82.87% for fastMNN, and 82.83% for Scanorama (Figure S7A). In the analyses where noise is spiked-in, mNN methods forced matching of almost 100% of cells with accuracy below 70% with increasing noise, whereas MARIO alerted the user of insufficient information for matching (Figure S7B). MARIO was over most resistant to cell-type composition changes (Figure S7C; error avoidance scores B cells: MARIO, 1.12; mNN methods, 0.46-0.96; NK cells: MARIO, 2.97; mNN methods, 0.55-1.03; neutrophils: MARIO, 2.08; mNN methods, 0.42-1.02; CD8 T cells: MARIO, 2.49; mNN methods, 0.65-1.17; CD4 T cells: MARIO, 1.65; mNN methods, 0.43-0.97; monocytes: MARIO, 1.61; mNN methods, 0.54-1.24).

### Accurate tissue architectural reconstruction reveals diverse lymphocyte populations

Inferring the spatial localization of biofeatures at the single-cell level is necessary for a holistic understanding of cellular processes *in situ* (22). Currently used multi-modal approaches to measure nucleic acids and proteins in their native tissue context are often limited by scale or resolution (9, 23, 25). We reasoned that a highly accurate cell matching and integration strategy, such as MARIO, could infer the spatial localization of transcripts within individual cells. We performed MARIO on spatially resolved data from murine splenic cells collected using antibody-based CODEX imaging (29 protein markers)(13) and data from dissociated murine splenic cells assayed using CITE-seq (206 protein markers) (46); 29 protein markers (all the markers in the CODEX dataset) were shared.

We first visually verified successful MARIO matching and integration using dimension-reduced t-SNE plots (Figure 4A). Cell-cell matching accuracy was high across all cell types: 87.69% for NK cells, 90.04% for neutrophils, 73.84% for macrophages, 83.72% for monocytes, 94.35% for DCs, 95.61% for CD8 T cells, 95.70% for CD4 T cells, and 93.99% for B cells (Figure S8A). This enabled highly accurate single-cell information transfer between cells measured using CITE-seq and CODEX spatially resolved cells (Figure 4B and Figure S8B). We visually observed highly concordant spatial organization of cell types annotated using CODEX or CITE-seq information and further observed a clear distribution pattern of transcripts corresponding to their expected spatial localization in the spleen (Figure 4B and Figure S8B). For example, *Il7r* is concentrated in the T cell zone as expected (47); *Myc* and *Cxcr5* are localized to activated and proliferating T and B cells within the germinal center (48, 49); *Ms4a1* and *Bhlhe41* are highly expressed in the B cell zone and B cells in the red pulp region (50–53); and *Il1b* is expressed outside the B cell zone (54). t-SNE overlays of the matched protein and RNA expression confirmed expected RNA expression profiles within given cell types (Figure S8C).

**Figure 4:**
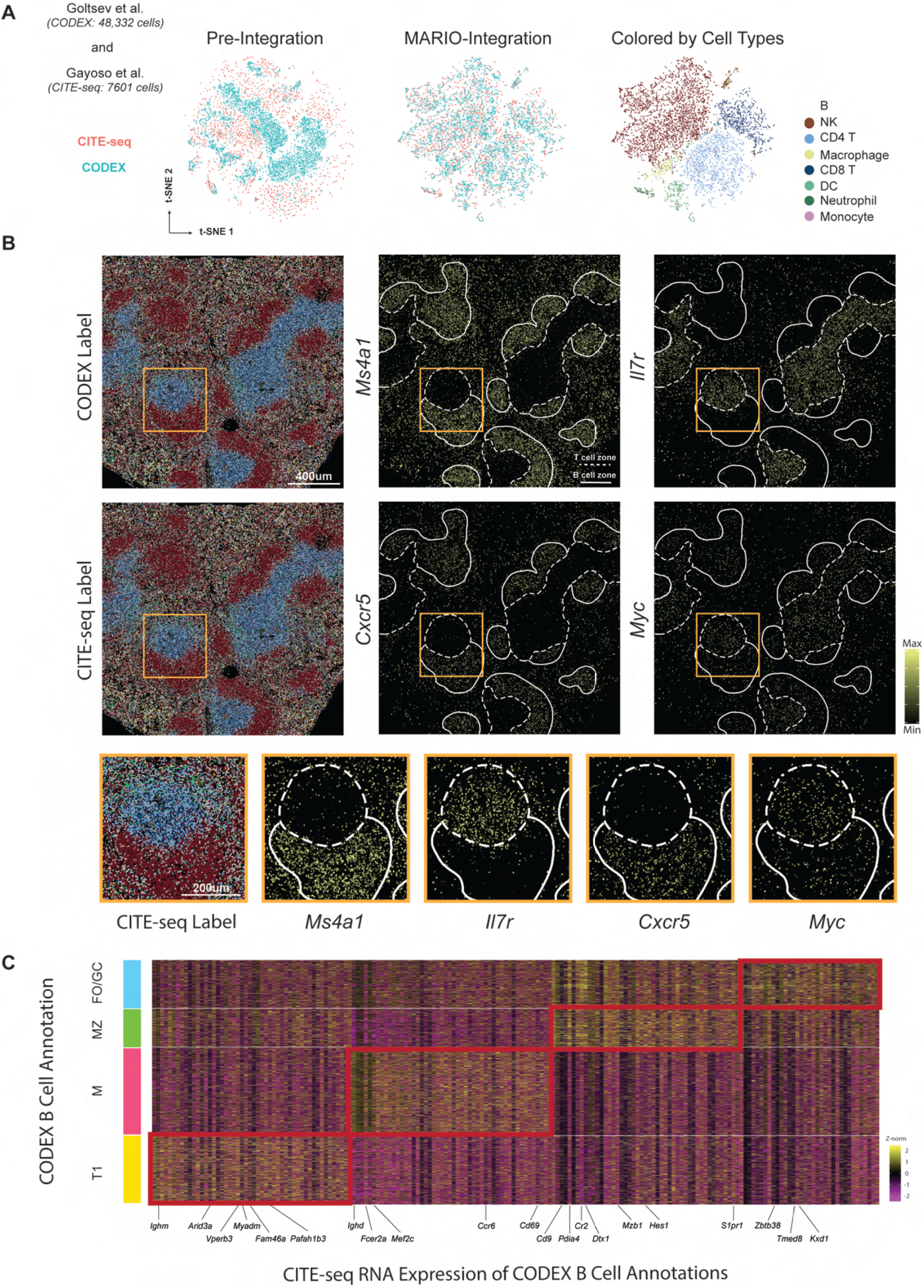
MARIO Integration of Suspension and Tissue Single-cell Measurements Enables Spatial Multi-omics. **(A)** t-SNE plots of murine spleen CITE-seq and CODEX cells, pre-integration and MARIO integration, colored by the dataset of origin (left and middle) or colored by cell type annotation (right). **(B)** A murine spleen section colored by the cell type annotation from CODEX (top left) or the label transferred annotation from CITE-seq (middle left). Examples of RNA transcripts ((*Il7r, Ms4a1, Cxcr5* and *Myc*) and their tissue-specific localization are inferred through MARIO integrative analysis (middle and right columns). An enlarged view of the tissue region demarcated by the orange box is shown in the bottom row. **(C)** Heatmap of differentially expressed genes (from matched CITE-seq cells) among subpopulations of CODEX B cells, gated based on CODEX proteins.

We next sought to further refine cells from the B lymphocyte lineage by gating the B cell population from the CODEX dataset based on B220, CD19, IgM, IgD, CD21/35, and MHCII. Four sub-populations of B cells were identified: Transitional type 1 B cells (T1), Marginal Zone B cells (MZ), Mature B cells (M) and Follicular/Germinal Center B cells (FO/GC) (Figure S8D). Visual inspection of the spatial location of these four subtypes of B cells confirmed localization within mouse spleens consistent with previous observations (Figure S8E) (55, 56). MARIO-matching thus enabled a detailed examination of the differentially expressed transcripts within these B cell subtypes resolved by CODEX, revealing a distinctive transcriptional program reflective of their phenotype (Figure 4C). For example, we observed signature landmark genes previously shown to demarcate these B cell subtypes from single-cell or bulk transcriptomic analysis (*Ighm, Arid3a*, and *Pafah1b3* for T1; *Ighd, Fcer2a*/*Cd23* and *Cd69* for M; *Cd9, Cr2* and *Mzb1* for MZ; *Zbtb38, Tmed8* and *Kxd1* for FO/GC)47 (57–59). These genes were significantly up-regulated (p-adjust < 0.05, Wilcoxon Test) in the corresponding gated populations of CODEX B cells.

For this CODEX to CITE-seq matching, MARIO had matching accuracy superior to mNN methods (Figure S9A). For the full 28-antibody shared panel, the total accuracy for MARIO was 87.76%, for Seurat it was 83.64%, for fastMNN it was 87.40%, and for Scanorama it was 82.70%. Dropping eight shared antibodies in the panel resulted in total accuracies of 85.31% for MARIO, 77.97% for Seurat, 82.01% for fastMNN, and 80.03% for Scanorama. MARIO prevented over-integration due to poor quality data, whereas the mNN methods forced matching (Figure S9B). MARIO was also robustness in resisting changes to cell-type composition (Figure S9C; error avoidance scores: DCs: MARIO, 1.63; mNN methods, 0.39-0.83; NK cells: MARIO, 1.66; mNN methods, 0.31-0.7; monocytes: MARIO, 1.82; mNN methods, 0.32-0.72; CD8 T cells: MARIO, 2.48; mNN methods, 0.53-1.23; CD4 T cells: MARIO, 2.24; mNN methods, 0.56-1.18; macrophages: MARIO, 1.77; mNN methods, 0.30-0.74).

### A COVID-19 lung molecular atlas reveals the role of complement activation in macrophages and related orchestrated immune responses

Single-cell profiling technologies have emerged as powerful tools in response to the ongoing COVID-19 pandemic. The deep functional characterization of clinical samples has provided critical insights into viral pathogenesis and tissue-specific host immune responses (60). Understanding these responses in their native tissue context has implicated potential therapeutic avenues (61, 62), but highly coordinated efforts are needed for an integrative understanding of the biological effects in COVID-19 (63).

We reasoned that the ability to perform integrative and inferential analysis across biological analogous clinical cohorts, measured at different institutions with varying technologies, would further our understanding of the facets of COVID-19 biology. We profiled 76 lung tissue regions from 23 individuals who succumbed to COVID-19 using CODEX high dimensional imaging with 50 markers, and MARIO-matched the macrophage population identified therein against those from bronchoalveolar lavage fluid (BALF) samples subject to CITE-seq with 250 surface markers (Figure 5A).

**Figure 5:**
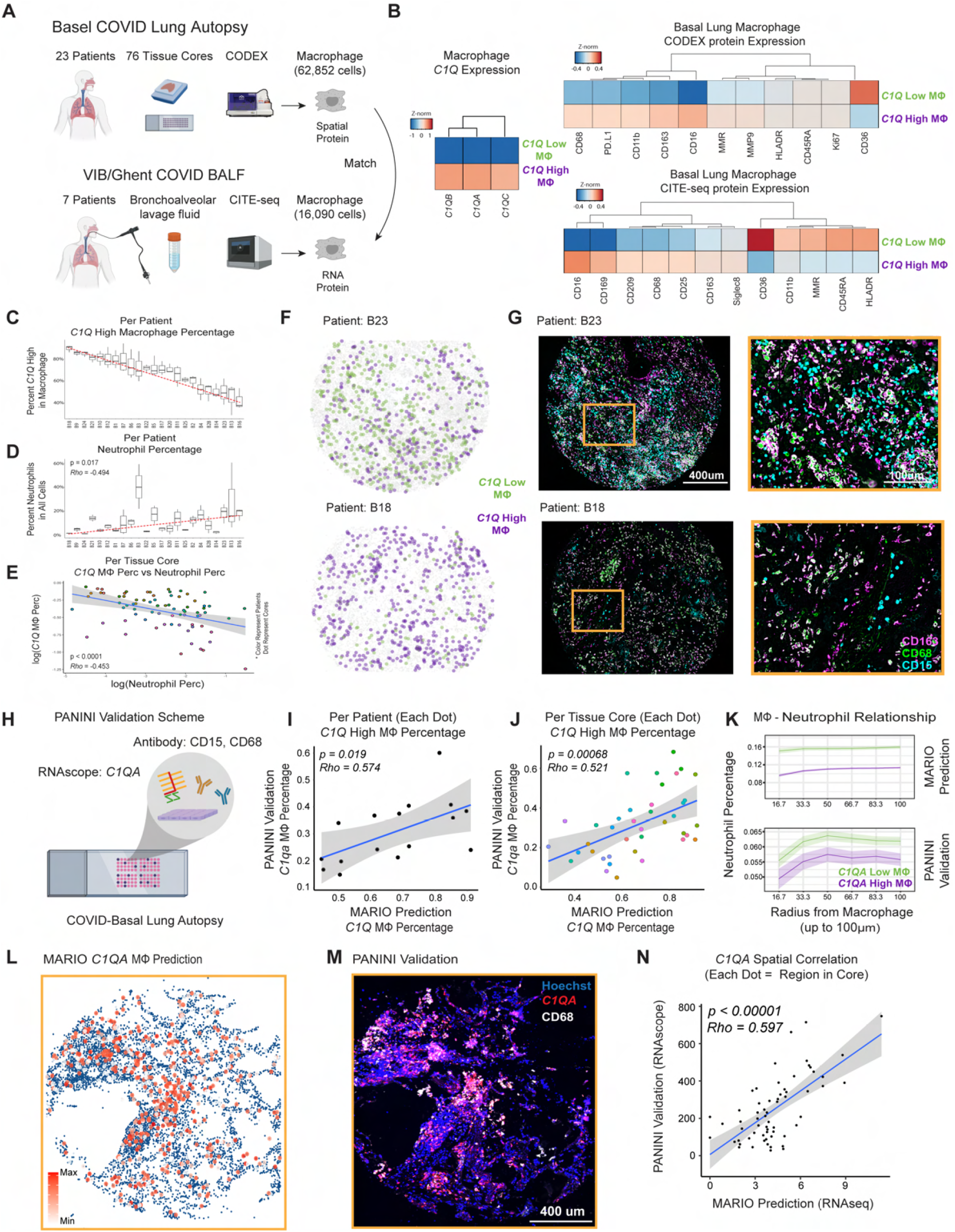
Integrative Spatial Multi-omic Analysis of Macrophages in COVID-19 patients with MARIO. **(A)** A schematic of the experimental and MARIO analysis on BALF and lung tissues from COVID-19 patients were measured from two independent studies via CITE-seq (from VIB/Ghent) and CODEX (University Hospital Basel/Stanford). Macrophages from the CODEX lung data were matched to those identified from BALF using CITE-seq using MARIO for integrative analysis. **(B)** Heatmaps of *C1Q* High and Low macrophages identified from CITE-seq, and their matched CITE-seq and CODEX expression patterns. **(C)** A ranked plot for macrophages from each patient in the CODEX data, as a percentage of *C1Q* High proportions. **(D)** Proportion of Neutrophils (as a percentage of all cell types) in each patient from the CODEX data, ranked by the same sequence as in (C). **(E)** A dot plot showing the relationship between *C1Q* High macrophages (Y axis) and Neutrophil percentage (X axis). Each dot represents a tissue core from the tissue microarray. **(F)** An representative pseudo image of two tissue cores colored with the locations of *C1Q* High and Low macrophages. **(G)** The CODEX multiplexed Images of the same two tissue cores in (F), with CD163, CD68 and CD15 antibody staining. An enlarged view of the region demarcated by the orange box is shown on the right. **(H)** An experimental schematic of PANINI to validate the spatial localization of *C1Q* macrophages on Basel/Stanford COVID-19 tissues. Slides were co-stained with probes detecting *C1QA* mRNA and antibodies targeting CD15 and CD68 proteins. **(I-J)** A dot plot showing the relationship between the proportion of *C1QA* High Macrophages (as a percentage of all macrophages) from the PANINI validation (Y axis) versus the MARIO prediction (X axis) per patient (I) or per tissue core (J). P-values and correlations were calculated using the Spearman-ranked test. **(K)** Anchor plots of Neutrophils as a function of distance from *C1QA* High (magenta) or *C1QA* Low macrophages (green) in MARIO predicted (above) or PANINI validated (below) experiments. **(L-M)** A representative tissue core with MARIO predicted *C1QA* expression levels in macrophages (left), and PANINI validated *C1QA* and CD68 signals (right). **(N)** Spatial-correlations between validation and prediction experiments were performed. The tissue core was divided into 10×10 regions, the summation of *C1QA* signals in macrophages were calculated and plotted for Mario and PANINI validation (P-value and correlation calculated by Spearman-ranked test).

We were able to stratify the macrophages into two populations based on their transcriptional signatures of complement pathway activity (Figure 5B; *C1Q* Low and *C1Q* High). Interestingly, we observed a positive correlation between the abundance of *C1Q* Low macrophages and patient body mass index (BMI; Figure S10D). Given that low serum *C1Q* levels have been reported in patients with severe COVID-19 (64), future studies should explore whether *C1Q* dysregulation can explain the positive association between obesity and risk for COVID-19-related hospitalization and death (65). The protein expression of these two classes of macrophages also partly corresponded to a M1 phenotype for *C1Q* Low macrophages, and an immunosuppressive M2 phenotype for *C1Q* High macrophages (Figure 5B). We further observed that the *C1Q* High transcriptional program was enriched in antigen processing and presentation, whereas that of the *C1Q* Low population consisted of several immune chemotaxis and migration pathways, including that of neutrophil chemoattractants (Figure S10A). The top differentially expressed transcripts included *CXCL8, CCL7* and *TMEM176B*, with previously described roles in regulating neutrophil recruitment and migration (66–68). The roles of proteins encoded by *IL1B, S100A8* and *CCL2* in the recruitment of aberrant neutrophils has been recently eluded in NHP and mice models of SARS-CoV-2 lung pathology (69), and are also reflected by elevated transcript levels in *C1Q* Low macrophages (Figure S10B).

In the five previously established functional clusters of interferon stimulated genes (ISG) (70, 71), we observed distinctive ISG transcriptional programs in *C1Q* Low and High macrophages (Figure S10C; p-adjust < 0.05, Wilcoxon Test) across all clusters (C1 & C2: RNA Process, C3: IFN Regulators - Antiviral effectors, C4: Metabolic Regulation, C5: Inflammation). Of particular interest is the C3 (Antiviral Activities) and C5 (Inflammation) clusters (Figure S10C; Green and Gold clusters). Our results suggest that in C1Q Low macrophages several previously described genes (including *SERPIN89, MX1, LGAPS3BP, SIGLEC1, CKAP4, CCL2* and *SPHK1*) that encode proteins reported to directly inhibit SARS-CoV-2 replication and entry are upregulated, but the failure to regulate and dampen this innate response paves the way to unchecked host immune responses and collateral tissue damage (72–76) (Figures S10C).

In line with the transcriptional signatures for aberrant neutrophil infiltration (Figure S10A), we noted a correlation between the presence of *C1Q* Low macrophages and increased infiltrating neutrophils (Figure 5C-E; Rho = −0.453, p < 0.0001). This elevated neutrophil presence was also confirmed visually (Figures 5F-G and S10E). Spatial cell-cell interaction analysis showed striking differences in these two subclasses of macrophages and their proximity with other cell types, such as high frequency of *C1Q* High macrophages to be proximal to CD4 and CD8 T cells, B cells, myeloid cells and other macrophages (Figure S10F). We next anchored *C1Q* High and Low macrophages for an anchor analysis (25) to understand the microenvironment as a function of distance around these two groups of macrophages. Our analysis confirmed the distinctive microenvironments and differences in immune orchestration around these macrophages, as evident from the differential organization of macrophages, plasma cells, vasculature and CD8 T cells (Figure S10G).

We finally performed Protein And Nucleic acid IN situ Imaging (PANINI)(25) to visualize the mRNA of a complement marker, *C1QA*, the neutrophil marker CD15 and the macrophage marker CD68 on COVID-19 tissue microarray sections to experimentally validate the spatially resolved gene expression patterns predicted by MARIO (Figure 5H). We confirmed the robust expression patterns of *C1QA* mRNA, CD68 and CD15 proteins in the tissue sections (Figure S10H). We observed a robust and significant correlation between the percentages of experimentally validated *C1Q* High macrophages and MARIO-predicted *C1Q* High macrophages percentage, both at the patient level (p = 0.019, Rho = 0.574) and at the per tissue core level (p = 0.000068, Rho = 0.521, Spearman Ranked test, Figures 5I and J). In line with anchor analysis from MARIO-inferred data, we confirmed a significantly decreased neutrophil density around *C1Q* High macrophages in the PANINI validation experiment (Figure 5K). The RNA spatial pattern from our PANINI experiment, performed on a separate, non-adjacent section of the same patient tissue core, recapitulated the prediction from the MARIO-matched data (Figure 5L and M). The spatial correlation between MARIO-predicted and PANINI-validated expression levels of *C1QA* in macrophages was highly consistent even between non-adjacent sections of the same tissue core (*C1QA* signal per region: p < 0.00001, Rho = 0.597, Spearman ranked test, Figure 5N). This rho value was close to the maximum possible spatial correlation of the tissue structure as determined using cell density per region (p < 0.00001, Rho = 0.602, Figure S10I), validating the highly accurate inferential capabilities of MARIO.

## Discussion

MARIO is a powerful matching and integration framework for single-cells that allows the retention of distinct features. It is thus particularly suitable for the integration of single-cell proteomic datasets with limited antibody panel overlap. We demonstrated that MARIO robustly and accurately matched cells across multiple sample types, assays, and species. Unlike current methodologies, MARIO performs pairwise matching of individual cells utilizing both shared and distinct features and is coupled with rigorous quality control steps. We benchmarked our algorithm across multiple datasets, and MARIO consistently outperformed other methods that were primarily designed for single-cell sequencing data and that are reliant upon the mNN matching algorithm. Importantly, MARIO inferential results allowed novel biologically interpretable insights. First, we demonstrated how CITE-seq data for human bone marrow cells could be leveraged to accurately delineate memory and naive T cell subtypes measured with a CyTOF panel lacking these naive/memory functional antibody markers. Second we showed that conserved and differential responses of human and NHP blood samples could be identified in data from different CyTOF experiments when matched using MARIO. Third, RNA transcripts could be spatially located within the murine spleen through the integration of CODEX and CITE-seq. Finally, two classes of complement pathway *C1Q* High and *C1Q* Low macrophages from COVID-19 BALF suspension cells analyzed by CITE-seq matched with COVID-19 lung autopsy CODEX data using MARIO delineated of the roles that these cell populations play in orchestrating immune responses to SARS-CoV-2 infection.

This MARIO analysis pipeline builds upon several novel and consolidated mathematical advances. First, the matching is constructed by globally (rather than locally) optimizing over a novel distance matrix that incorporates both the explicit correlations in shared features and the hidden correlations among distinct features. Second, the accuracy and robustness of the matching is ensured by two theoretically principled quality control processes, the Matchability Test and the Jointly Regularized Filtering (77). Third, the integrated embeddings are obtained via CCA or gCCA which incorporates the information in both the shared and distinct features.

In spite of the clear advantages of MARIO, it has some technical limitations. First, the accuracy and robustness come at the cost of longer analysis times compared to mNN-based approaches. Given the globally optimal nature of the core matching algorithm implemented in MARIO, the time required to run the MARIO pipeline is cubically related to the number of cells; in contrast, time required for mNN-based methods is quadratically related to the number of cells. To circumvent this, we developed a sparsification technique that reduces the search space, which accelerates the matching process. Empirically, we found that MARIO can be run on datasets with moderate sample sizes within reasonable time frames: The execution time for 50,000 cells took 10 minutes, with a peak memory usage of approximately 7 GB (Figure S11). Second, although MARIO out performs mNN-based methods in the scarce shared feature regime, its success relies on the existence of shared features. This may not be the case in certain scenarios such as when integrating RNA-only and protein-only data. Future work incorporating methods that enable inference of protein levels from transcript levels will no doubt allow methods such as MARIO to have even broader applicability.

The need to study biological processes within their tissue context is increasingly evident, with direct relevance to the physiological context of health and disease. Simultaneous single-cell measurement of nucleic acids and proteins in their spatial context remains challenging, despite recent advancements (25, 26, 78), and it remains limited by factors including resolution and requirements for tissue fixation. The ability to match similar biological samples measured using distinctive single-cell assays will be paramount for hypothesis generation and guidance for experimental design. We are confident that MARIO will serve as a useful methodology and resource for the community with direct applications to a plethora of experimental platforms and biological contexts.

## Materials & Methods

### Cell matching

Suppose we have two datasets *X* and *Y*, where 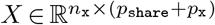 consists of *n*_x_ cells and (*p*_share_ + *p*_x_) features and 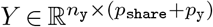 consists of *n*_y_ cells and (*p*_share_ + *p*_y_) features. Without loss of generality, we assume *n*_x_ ≤ *n*_y_. Among all the features, *n*_share_ features are shared across both datasets, whereas the rest of the features are distinct to either *X* or *Y*. Thus, we can write both datasets as horizontal concatenations of a shared part and a distinct part:

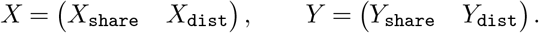

The *cell matching* between *X* and *Y* is defined as an injective map Π, represented as a binary matrix of dimension *n*_x_ × *n*_y_, such that Π_*i,i*_*′*= 1 if and only if the *i*-th cell in X share a similar biological state with the *i*^′^-th cell in *Y*.

#### Initial matching with shared features

We first construct an initial estimator of Π using shared features alone. The procedure starts by denoising the shared parts via thresholding their singular values. Consider the singular value decomposition of the vertical concatenation of *X*_share_ and *Y*_share_:

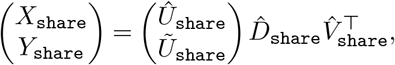

where the vertical concatenation of 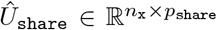 and 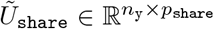 collects the left singular vectors, 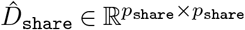 is a diagonal matrix that collects the singular values in descending order, and 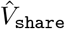 collects the right singular vectors. Let 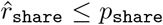 be the number of components to keep. In the MARIO package, we denote 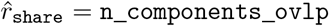. We then compute the denoised version of *X*_share_ and *Y*_share_ by

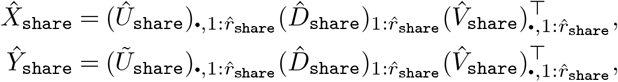

respectively, where for a matrix *A*, we let *A*_•,1:*r*_ denote its first *r* columns and for a diagonal matrix *D*, we let *D*_1:*r*_ denote the submatrix formed by taking its first *r* rows and columns. We then construct a cross-data distance matrix 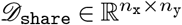, whose entries are given by

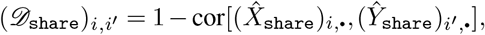

where 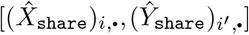 is the Pearson correlation coefficient between the *i*-th row of 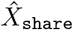 and the *i*^′^-th row of *Ŷ*_share_. The initial estimator of Π is given by the solution of the following optimization problem:

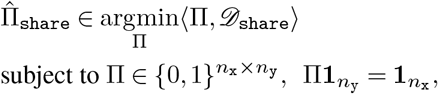

where for two matrices *A* and *B*, we let ⟨*A, B*⟩ = ∑_*i,i*_*′ A*_*i,i*_*′ B*_*i,i*_*′* denote the Frobenius inner product. This optimization problem is an instance of minimal weight bipartite matching (a.k.a. rectangular linear assignment problem) in the literature (79).

#### Refined matching with distinct features

Given the initial matching 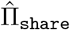, we can approximately align cells in *X* and *Y* : the rows of *X* and 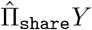 correspond to pairs of cells with similar biological states, up to a certain level of mismatches induced by the estimation error of 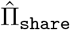. Despite mismatches, such an approximate alignment opens up the possibility of estimating the latent representations of *X* and *Y* by CCA.

Assuming both *X* and *Y* are standardized so that their columns are centered and scaled to have unit standard deviation. Then their empirical covariance and cross-covariance matrices are given by

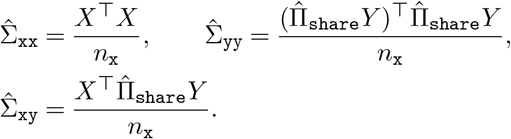

The first pair of sample canonical coefficient vectors is given by

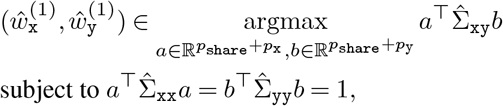

and the first sample canonical correlation is given by 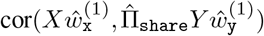. Now, for 2 ≤ *j* ≤ *p*_share_ + min(*p*_x_, *p*_y_), the *j*-th pair of sample canonical coefficient vectors is successively defined as

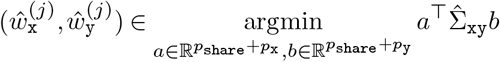

subject to *a*^T^Σ_xx_*a* = *b*^T^Σ_yy_*b* = 1,

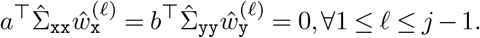

In parallel, the *j*-th sample canonical correlation is given by 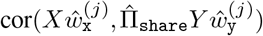. Let 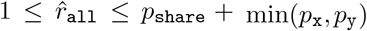 be the number of components to keep. In the MARIO package, we denote *r*_all_ = n_`components`_`all`. Collecting top 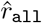 sample canonical vectors into matrices

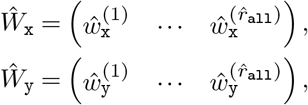

the latent representation of *X* can be estimated by *XŴ*_x_, the sample canonical scores of *X*. That is, we use *Ŵ*_x_ to project *X* onto the latent space. The same projection can be done on *Y* data by computing *Y Ŵ*_y_, so that the resulting matrix approximately lies in the same latent space as *XŴ*_x_.

To this end, we compute the cross-data distance matrix *𝒟*_all_ directly on the latent space, whose entries are given by

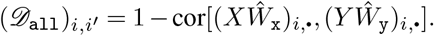

We finally solve for a refined matching by

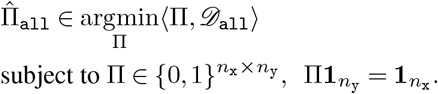

#### Interpolation of initial and refined matchings

The quality of the refined matching 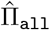 is highly contingent upon the quality of the distinct features. If the distinct features are extremely noisy, incorporation of them may hurt the performance, in which case it is more desirable to revert back to the initial matching 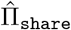. We develop an data-adaptive way of deciding how much distinct information shall be incorporated when we estimate the matching from the data.

To start with, we cut the unit interval [0, 1] into grids (e.g., {0, 0.1,…, 0.9, 1}). For each *λ* on the grid, we interpolate the two kinds of distance matrices by taking their convex combination

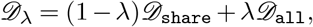

from which we can solve for the *λ*-interpolated matching

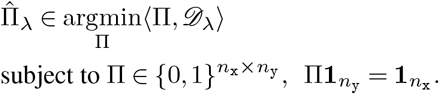

Note that 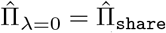 and 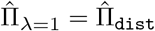. After aligning *X* and *Y* using 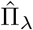, we compute top k sample canonical correlations (in the MARIO package denoted as `top_k`, and defaulted to 10), whose mean is taken to be a proxy of the quality of 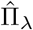. We then select the best 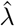 according to this quality measure and use 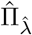 afterwards.

### Quality control

#### Test of matchability

In extreme cases, the two datasets *X* and *Y* may not have any correlation at all, and thus any attempt to integrate both datasets would give unreliable results. For example, some methods, when applied to uncorrelated datasets, would pick up the spurious correlations and hence resulting in over-integration. A robust procedure should be able to tell and warn the users when the resulting matching estimator might be of low quality. We develop a rigorous hypothesis test, termed matchability test, for this purpose.

The matchability test starts by repeatedly drawing *B* i.i.d. copies of *n*_x_-dimensional (potentially asymmetric) Rademacher random vectors 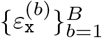 and another *B* i.i.d. copies of *n*_y_-dimensional Rademacher random vectors 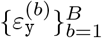. That is, for each 1 ≤ *b* ≤ *B*, we have 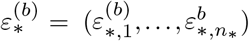, and 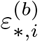 is + 1 with probability 1 − *p*_flip_ and is − 1 otherwise for any 1 ≤*i* ≤ *n*_*_, where * is the placeholder for either `x` or `y`. The parameter *p*_flip_ (denoted as `flip_prob` in MARIO package and defaulted to 0.2) controls the “sensitivity” of the test — a lower value of *p*_flip_ means that a more accurate matching is needed to pass the matchability test. For every *b*, we generate a fake pair of datasets by flipping the signs of each row of *X* and *Y* :

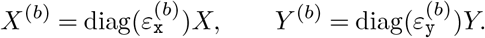

After such a sign-flipping procedure, the majority of the correlation (i.e., the inter-dataset covariance structure) between *X* and *Y*, if exists, is destroyed. On the other hand, the intra dataset covariance structures of both *X* and *Y* are preserved.

As a result, if we run any matching algorithm with *X*^(*b*)^ and *Y* ^(*b*)^ as the input, the resulting estimator 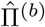 would be of low quality, in the sense that if we align *X*^(*b*)^, *Y* ^(*b*)^ using 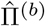 and run CCA, the resulting sample canonical correlations will be small. In our implementation, we calculate the mean of `top_k`, and defaulted to 10), which we denote as 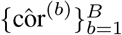.

The matchability test proceeds by running the same algorithm on the real datasets *X, Y*, aligning them using the estimator 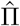, and calculate the mean of `top_k` sample canonical correlations, which we denote as côr. The final *p*-value for testing the null that *X* and *Y* are uncorrelated is given by the proportion of 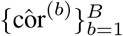 that are larger than the observed côr.

#### Jointly regularized filtering of low-quality matched pairs

Even if the two datasets *X* and *Y* are highly correlated (and thus the matchability test gives a small *p*-value), the estimated matching 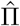 might still be error-prone. This could happen, for example, when certain cell types exist in *X* but are completely absent in *Y*. We develop an algorithm that automatically filters out the low-quality matched pairs in 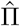.

Assume there are *K* cell types present in either *X* or *Y*. In the MARIO package, we denote *K* = n_clusters (default = 10). Let 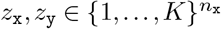 be the unknown ground truth cell type labels of *X* and 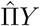, respectively. The fact that *X* and *Y* have passed the matchability test tells that *z*_x_ and *z*_y_ should agree on most coordinates. However, it is entirely possible that there exists a sparse subset of {1, …, *n*_x_} on which *z*_x_ and *z*_y_ disagree, and our goal is to detect this sparse subset and disregard them in downstream analyses. To achieve this goal, we consider the following regularized *k*-means objective:

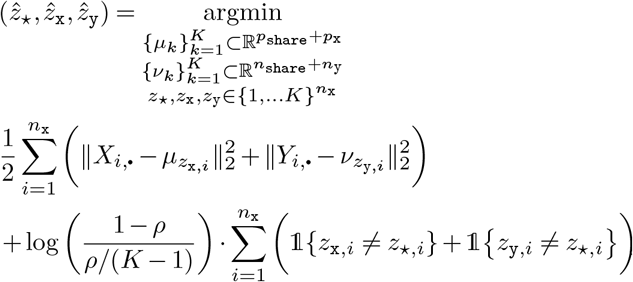

where ‖ · ‖_2_ is the *𝓁*_2_ norm and 𝕝 {·} is the indicator function. The above objective function is a superposition of two parts. The first part is the classical *k*-means objective for *X* and *Y*, and the second part is a regularization term that imposes penalties when the estimated *X*-label 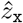 and *Y*-label 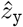 are too far-away from a “global” label 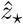.

After solving the above objective function, if 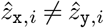, then there is evidence that the matched pair 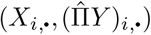 is spurious, and is thus disregarded in the downstream analyses. The parameter *ρ* controls the strength of regularization: if *ρ* = 1 − 1*/K*, then there is no regularization at all, whereas if *ρ* = 0, we effectively require 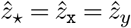. Thus, we can naturally control the “intensity” of such a filtering procedure by choosing a suitable *ρ*. In fact, under a hierarchical Bayesian model, the parameter *ρ* has a rather intuitive interpretation as the probability of disagreement between *z*_*⋆,i*_ and *z*_x,*i*_ (or between *z*_*⋆,i*_ and *z*_y,*i*_) (77). If the model is correctly specified, then the expected proportion that should be filtered out is given by 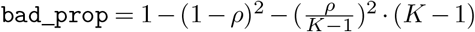.

We solve the regularized *k*-means objective via a warm-started block coordinate descent algorithm. The algorithm starts by computing initial estimators 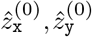 of *z*_*x*_, *z*_*y*_ via spectral clustering (80): we compute the sample canonical scores of *X* and 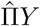, average them, and apply the classical *k*-means clustering on top *K* eigenvectors of the averaged score to get 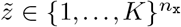. We then let 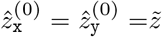. The number of canonical scores to keep is denoted as `n_components_filter` in the MAIRO package (default = 10).

Suppose at iteration *t*, the current estimators of *z*_x_, *z*_y_ are given by 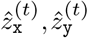, respectively. We run block coordinate descent as follows:

1. Given 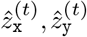, the current estimators of {*μ*_*k*_}, {*v*_*k*_} are given by

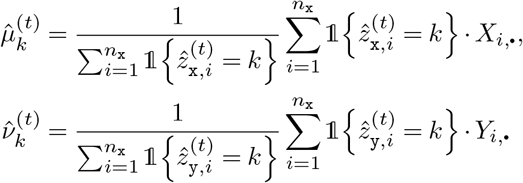

for any 1 ≤ *k* ≤ *K*.
2. Given 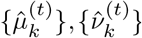, the next estimators of *z*_*\**_, *z*_x_, *z*_y_ are given by

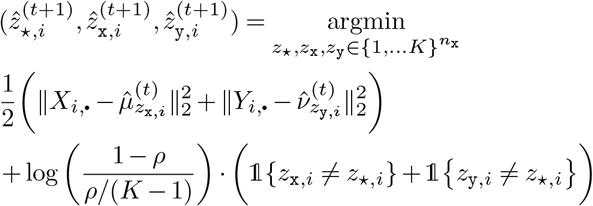

for any 1 ≤ *i* ≤ *n*_x_. The above problem is solved via a careful enumeration procedure. We first hypothesize that 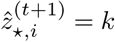 for some 1 ≤ *k* ≤ *K*. Given this hypothesis, we can solve for the best 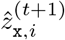 by enumerating all *K* possible choices of labels. The same thing can be done to solve for the best 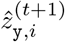. Hence, we can compute the best value of the above objective function under the hypothesis that 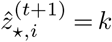. We can then solve for the global optimal 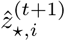 by enumerating and comparing the objective values under every possible hypothesized value of 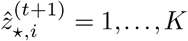. Given the global optimal 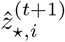, the global optimal 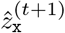 and 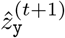 can be easily extracted.

In our implementation, we run the above block coordinate descent procedure for 20 iterations.

### Downstream analysis after cell matching

#### Joint embedding

After running jointly regularized filtering on the best interpolated estimator 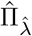, we get a pair of aligned datasets 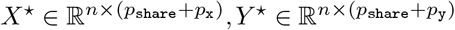, whose rows correspond to cells of similar types and *n* is the number of remaining cell-cell pairs after filtering. Then, we run CCA on *X*^*⋆*^, *Y* ^*⋆*^ and collect the first n pairs of sample canonical scores (scaled within dataset) as the final embeddings. Since the rows of *X*^*⋆*^ and *Y* ^*⋆*^ are approximately aligned, other standard methods for joint embedding (e.g., partial least squares) can also be applied.

#### Label transfer via k-NN matching

The interpolated distance 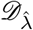 can be used to do label transfer via *k*-nearest-neighbors. Suppose we know the cell type labels for all cells in *Y* but the corresponding labels for cells in *X* is missing. Then for the *i*-th cell in *X*, we can predict its label by finding the *k*-nearest cells (we denote *k* = knn in the MARIO package) in *Y* according to 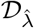 and taking the majority vote.

### Extensions

#### Matching more than two datasets

Suppose we have *L* datasets 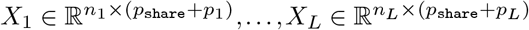. For 2 ≤ *𝓁* ≤ *L*, we run the usual two-dataset procedure to estimate the matching between cells in *X*_1_ and cells in *X*_*𝓁*_ by 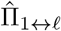. We then run jointly regularized filtering on each 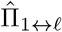 separately and keep the cells in *X*_1_ that survive all *L* − 1 rounds of filtering. This gives us a cell-to-cell matching among the *L* datasets, from which we can construct row-wise aligned datasets 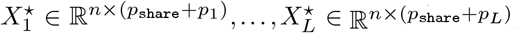, where *n* is the number cells in *X*_1_ that survived all *L* − 1 rounds of filtering.

To jointly embed all the aligned datasets, we use generalized canonical correlation analysis (gCCA) (81). It is well known that gCCA does not admit a unique formulation (82). We take the following formulation which best suits our goal of obtaining joint embeddings:

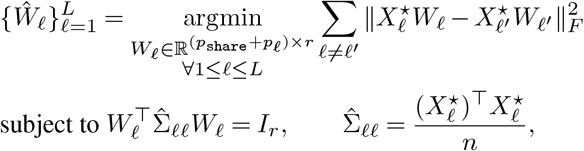

where ‖ · ‖_*F*_ is the Frobenius norm, 1 ≤ *r* ≤ *p*_share_ +min_*𝓁*_ *p*_*𝓁*_ is the number of components to keep, and *X*_*𝓁*_ *Ŵ*_*𝓁*_ is the embedding for the *𝓁*-th dataset.

To solve the above optimization problem, we take a block coordinate descent approach. This approach again needs preliminary estimators 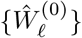. To obtain those preliminary estimators, we first run the classical CCA on the first two datasets and obtain the projection matrices 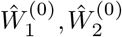, so that 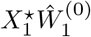 and 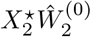 are the sample canonical scores for 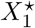 and 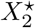, respectively. Then, for each *𝓁* ≥ 3, we run least squares regression using 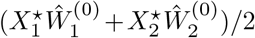 as the response and 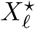 as the feature matrix. The resulting regression coefficient is then taken to be 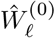.

Given the preliminary estimators, we are ready to enter the block coordinate descent iteration. We first demonstrate how to solve for the first columns of {*Ŵ*_*𝓁*_}. Suppose at iteration *t*, we are given preliminary estimators 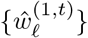, where 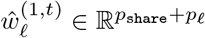. We then proceed as follows. For every 1 ≤ *𝓁* ≤ *m*, we run a least squares regression with the response being the current average scores (not counting *𝓁* itself), i.e., 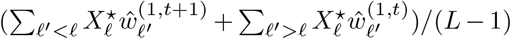, and with the feature matrix being 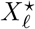. Denote the resulting regression coefficient as 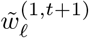. We take 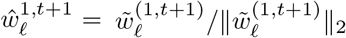. We run the above procedure for 500 iterations and let 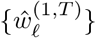 be the first columns of {*Ŵ*_*𝓁*_}.

We now discuss how to solve for the *j*-th columns of {*Ŵ*_*𝓁*_}, where *j* ≥ 2. We start by running a least squares regression with 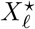 (i.e., 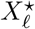, being the response and the first *j* − 1 scores of 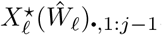, where (*Ŵ* _*𝓁*_)_· 1: j − 1_ is the first *j*− 1 columns of *Ŵ*_*𝓁*_) being the feature matrix. The residual of this regression is denoted as 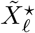. Now suppose at iteration *t*, we are given preliminary estimators 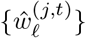, where 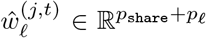. We proceed as follows. For every 1 ≤ *𝓁* ≤ *L*, we run a least squares regression with the response being 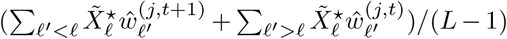, and with the feature matrix being 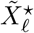. Denote the resulting regression coefficient as 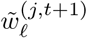. We then run a least squares regression with the response being 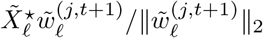 and the feature matrix being 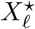. The resulting regression coefficient is taken to be 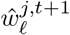. We run the above procedure for 500 iterations and let 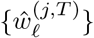 be the *j*-th columns of {*Ŵ*_*𝓁*_}.

#### Speeding up cell matching via distance sparsification

Standard implementations of the one-to-one matching run in 𝒪 ((*n*_x_ + *n*_y_)^3^) time. However, if the distance matrix *𝒟* is sparse (i.e., a lot of entries are infinity, meaning that such a pair is a priori infeasible), then the time complexity can further be reduced. For example, if one regards the distance matrix as a bipartite graph and let (*i, j*) denote an edge if *D*_*ij*_ *<* ∞, then it is possible to solve the problem in 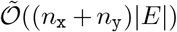 time, where *E* is the number of edges and 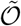 hides poly-log factors (83).

A natural attempt is to manually sparsify *𝒟* so that for each row, only *k ≪ n*_y_ smallest entries are finite. Let *𝒟* ^(*k*)^ be the sparsified matrix. In theory, there exists a critical value of *k*^*⋆*^ such that: (1) the distance matrix 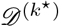 can give a valid matching; and (2) if one sparsifies it further (i.e., use *D* ^(*k*)^ for *k < k*^*⋆*^), then there is no valid matching. We give an algorithm for computing this critical value. For any fixed *k*, we can test if *𝒟* ^(*k*)^ can give a valid matching by computing the maximum-cardinality matching, which can be done in 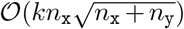 time using the Hopcroft–Karp algorithm (84). We can then use binary search to search for the critical value *k*^*⋆*^ In the worst case (i.e., when *k*^*⋆*^ = *n*_y_), the whole procedure runs in 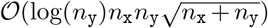 time, which is already much faster than the 𝒪 ((*n*_x_ + *n*_y_)^3^) time needed to compute the matching using the original distance matrix. In practice, since *k*^*⋆*^ is usually very small compared to *n*_y_, the running time of the whole procedure can be even faster. This procedure generalizes the strategy taken by (85), which only works when the distance matrix is computed using a single feature.

Given the knowledge of *k*^*⋆*^, we sparsify the distance matrix with some user-specified *k* ≥ *k*^*⋆*^ (denoted as `sparsity` in the MARIO package) and apply the LAPJVsp algorithm (an algorithm specifically designed to tackle sparse inputs) (86) to compute the matching.

In practice, we can further speed up the matching process by randomly splitting the data into *n* (in MARIO package denoted as `n_batch`) evenly-sized batches, computing the matching for each batch, and stitching the batch-wise matchings together.

### Details on data pre-processing and analysis

#### Code and data availability

MARIO and related tutorials are freely available to the public at GitHub: https://github.com/shuxiaoc/mario-py. Data and Code to regenerate the main and supplementary figures are also deposited to GitHub.

#### Preprocessing and analysis of human bone marrow datasets

CyTOF data measuring 32 proteins in healthy human bone marrow cells from levine et al (32)) was downloaded from GitHub https://github.com/lmweber/benchmark-data-Levine-32-dim.

Cells gated as HSPCs, CD4 T cell, CD8 T cell, B cell, monocyte, NK cell and pDC from the paper were selected and a total of 102,977 cells were used. CITE-seq dataset measuring 25 proteins and RNA expression of healthy human bone marrow cells was acquired using `bmcite` in the R package `SeuratData`. Cells annotated as HSPCs, CD4 T cell, CD8 T cell, B cell, monocyte, NK cell, and pDC from the paper, comprising a total of 29,007 cells, were used. During matching, CITE-seq cells were used to match against CyTOF cells, where the input of CITE-seq cells were prenormalized counts from `bmcite` and the input of CyTOF cells were values with arcsine transformation (cofactor = 5). The MARIO parameters used are `n_components_ovlp` = 10, `n_components_all` = 20, `sparsity` = 1000, `bad_prop` = 0.2, and `n_batch` = 4.

t-SNE plots were generated using the scaled shared protein features across datasets (pre-integration) or the first 10 components for the CCA scores (MARIO integration), using the `Rtsne()` function with default settings in R package `Rtsne`. The heatmap was produced using `heatmap.2()` in the R package `gplots`, with z-scaled CITE-seq and CyTOF protein expression levels. The matched or original values of protein/RNA overlaid with t-SNE plots were generated with the function `Featureplot()` in R package `Seurat`. The detailed process of benchmarking MARIO against other methods is further described in the Benchmarking section in the Supplementary Methods section.

#### Preprocessing and analysis of cross species H1N1/IFN gamma challenged datasets

CyTOF data measuring 42 proteins in blood cells from humans challenged with H1N1 (40) virus was acquired from flow repository FR-FCM-Z2NZ 39. Three donors were used (id = “101”, “107”, “108”). The dataset was randomly downsampled to 120,000 cells, arcsine transformed with cofactor = 5, and subsequently clustered via the default `Seurat` clustering pipeline with all available antibody markers. Cell types were then manually annotated based on their expression profile. A total of 102,147 annotated cells were used. CyTOF data measuring 39 proteins of whole blood cells from human, rhesus macaque and cynomolgus monkey challenged with Interferon gamma (37) were acquired from flow repository FRFCM-Z2ZY 35. Three donors of each species (human: “7826”, “7718”, “2810”; rhesus macaque: “D00522”, “D06022”, “D06122”; cynomolgus monkey: “D07282”, “D07292”, “D07322”) were used. Cells gated as Erythrocytes, Platelets and CD4+CD8+ cells in the paper were excluded from downstream analysis. Each individual dataset was randomly down-sampled to 120,000 cells, arcsine transformed with cofactor = 5, then clustered with `Seurat` using all the markers, followed by manually annotation and then removal of cells with ambiguous annotations. Total cell numbers for matching were 114,175 (human); 112,218 (rhesus macaque); 91,409 (cynomolgus monkey). During matching, human H1N1 challenged cells were matched against human, rhesus macaque and cynomolgus monkey IFN gamma-stimulated cells separately, and cells that matched across all four datasets were used for downstream analysis. The MARIO parameters used are `n_components_ovlp` = 20, `n_components_all` = 15, `sparsity` = 1000, `bad_prop` = 0.1, and `n_batch` = 4.

The t-SNE plot was produced by the scaled shared protein features across the dataset (pre-integration) or the first 10 components of the generalized CCA scores (MARIO integration), using the `Rtsne()` function with default setting in R package `Rtsne`. For visualization purposes, cell numbers were downsampled to 20,000 each dataset (80,000 cells in total) for t-SNE visualization. Euclidean distances between matched cells were calculated based on the integrated generalized CCA scores. Accuracy for MARIO matching results among cell types was generated by 5 repeated measurements on a randomly subsampled 5000 matched cells, and the balanced accuracy was calculated with the function `confusionMatrix()` in the R package `caret`. The expression level of Ki-67, pSTAT1 and p38 overlaid on each individual dataset’s t-SNE plots was produced with the function `Featureplot()` in R package `Seurat`. Violin plots were produced based on normalized (`scale()` function, within each dataset) values of Ki-67, pSTAT1, and p38 for Monocytes, CD4 T cells subpopulations with `ggplot2`.

#### Preprocessing and analysis of murine spleen datasets

Tiff files of CODEX multiplexed imaging data for BALBc mouse spleen, with 29 antibodies, were acquired (13) (sample ID: ‘balbc-1’). Segmentation was performed with a local implementation of Mesmer (87), with weights downloaded from: https://deepcell-data.s3-us-west-1. amazonaws.com/model-weights/Multiplex_ Segmentation_20200908_2_head.h5. Inputs of segmentation were DRAQ5 (nuclear) and CD45 (membrane). Signals from the images were capped at 99.7th percentile, with prediction parameter `model_mpp` = 0.8. Lateral spillover signals were cleaned using REDSEA (88) with the whole cell compensation flag as previously described. To clean out aggregated B220 signals in the dataset, B220 signal inside the cytoplasm (defined by 7 pixels towards the inside of the cell boundary), was removed. Afterwards, cells with DRAQ5 signal value less than 80 were removed and signals were scaled to 0-1, with percentile cutoffs of 0.5% (floor) and 99.5% (ceiling). Cells were subsequently clustered via `Seurat`, using CODEX markers: CD45, Ly6C, TCR, Ly6G, CD19, CD169, CD3, CD8a, F480, CD11c, CD27, CD31, CD4, IgM, B220, ERTR7, MHCII, CD35, CD2135, NKp46, CD1632, CD90, CD5, CD79b, IgD, CD11b, CD106. Another round of sub-clustering was then performed for dendritic cells, and macrophage populations before manual annotation of clusters. A total of 48,332 cells labeled as B cell, CD4 T cell, CD8 T cell, Dendritic cell, Macrophage, Monocyte, Neutrophil, and NK cells were used for MARIO matching. CITE-seq data 45 of murine spleen/lymph node samples from a panel of 206 antibodies were downloaded from GitHub: https://github.com/YosefLab/totalVI_reproducibility/tree/master/data. Only B, CD4 T cell, CD8 T cell, dendritic, macrophage, neutrophil, and NK cells originating from the spleen, a total of 7601 cells, were used. For matching, the input of CODEX cells are post-compensated, aggregation corrected values, excluding the Ter119 red blood cell channel. CITE-seq input were the downloaded raw counts. The CITE-seq dataset was duplicated to improve the matchability, and CODEX cells subsequently matched against CITE-seq cells, with MARIO parameters: `n_components_ovlp` = 20, `n_components_all` = 15, `sparsity` = 1000, `bad_prop` = 0.05, `n_batch` = 32, `knn` = 15.

The t-SNE plots were produced using the scaled, shared protein features across datasets (pre-integration) or the first 10 components for the CCA scores (MARIO integration), using the `Rtsne()` function with default settings in R package `Rtsne`. For visualization purposes, both datasets were downsampled to 8000 matched cells from each modality (16,000 cells in total) for t-SNE plotting. Pseudo-images of the CODEX murine spleen were colored by their cell-type annotations (Cell type based on CODEX protein annotation; Label transfering from CITE-seq annotation) and matched RNA expression levels. The label transfer of CITE-seq annotation shown in the figure was done using *k*-NN (*k* = 15) on the MARIO distance matrix, to ensure all CODEX cells have an annotation. The RNA expression value for pseudo-imaging plotting was capped to the 80% percentile (values equal to 0 were omitted) of that gene. For gating of B cell subtypes, CODEX proteins B220, CD19, IgM, IgD, CD21/35 and MHCII were used, and manually gated in cellengine https://cellengine.com/. Heatmaps of matched RNA expression level of CODEX B cell subpopulations was produced via the function `DoHeatmap()` in the R package `Seurat`, with top 50 differentially expressed genes identified in each subpopulation, via the function `FindAllMarkers()` in `Seurat`.

#### COVID-19 human tissue specimen collection

Lung tissues from patients who succumbed to COVID-19 were obtained during autopsy at the University Hospital Basel, Switzerland. Tissues were processed as previously described (89) and collection was approved by the ethics commission of Northern Switzerland (EKNZ; study ID #2020-00969). All patients or their relatives consented to the use of tissue for research purposes. Tissue microarrays were generated from these tissue samples in-house at the University Hospital Basel, Switzerland.

#### Preprocessing and analysis of COVID patient macrophage datasets

CODEX on COVID-19 samples from University Hospital Basel: CODEX acquisition of the COVID-19 tissue microarrays were performed, and post-processing and cell type annotation executed as previously described (90, 91). Data from 23 COVID-19 patients (76 tissue cores; manuscript in preparation) were acquired, and a total of 62,852 macrophages that were annotated were used for MARIO matching. Processed counts of CITE-seq data acquired with a panel of 250 antibodies from bronchoalveolar lavage fluid washes from COVID-19 patients (VIB/Ghent University Hospital) was acquired from COVID-19 Cell Atlas59. Cells from 7 COVID-19 patients (COV002; COV013; COV015; COV024; COV034; COV036; COV037) were selected, clustered, and manually annotated on a per patient level based on their protein features, using `Seurat` as previously described. A total of 16,090 macrophages were annotated and used for subsequent MARIO matching. During MARIO matching, CODEX macrophages were matched against CITE-seq macrophages, with the MARIO running parameters: `n_components_ovlp` = 25, `n_components_all` = 25, `sparsity` = 1000, `bad_prop` = 0.1, and `n_batch` = 20.

CODEX macrophages were clustered based on their matched *C1Q* mRNA expression levels (*C1QA, C1QB* and *C1QC*) using the function `hcut()` with k = 2 and stand = TRUE in the R package `factoextra`. Heatmaps were produced with the scaled values from CITE-seq or CODEX, via function `heatmap.2()` in R package `gplots`. Cell-cell interaction and binned anchor analysis were performed as previously described 25. In brief, for each individual *C1Q* High or Low macrophage, the Delaunay triangulation for neighboring cells (within 100μm) was calculated based on the XY position with the `deldir` R package. To establish a baseline distribution of the distances, cells were randomly assigned to existing XY positions, for 1000 permutations. The baseline distribution of the distance was then compared to the observed distances using a Wilcoxon test. The log2 fold enrichment of observed mean over expected mean for each interaction type was plotted for interactions with a p-value < 0.05. For the binned anchor analysis of *C1Q* High or Low macrophages, all cells within a 100μm range were extracted and the average percentage of specific cell types in each radius bin (in 16.66um increments) were calculated and plotted. Differential expression gene analysis was performed using the function `FindMarkers()` in the R package `Seurat`. The violin plot of DE genes were created with `ggplot2`, where mRNA expression values were normalized between 0-1 for visualization purposes. GO term analysis was conducted via the Gene Ontology tool (92, 93) (with the biological process option activated), with the input as lists of genes that were either significantly upregulated in *C1Q* High or Low macrophages. Heatmaps of the expression pattern of differentially expressed ISG genes (identified via `FindMarkers()`), filtered using a list of 628 ISGs with functional annotations 67 in macrophages, was plotted with the function `heatmap.2()` from the R package `gplots`. Correlations between *C1QA* macrophage percentages and neutrophil percentages were calculated with the R function `cor()` with method `spearman`.

#### PANINI Validation with COVID-19 Lung Tissue Samples

Protease-free combined ISH + antibody validation experiments using PANINI as previously described (25). In brief, TMA cores cut onto glass coverslips were baked at 70°C for 1hr and then transferred to 2 × 5 min xylene washes, followed by deparaffinization steps 2 × 100% EtOH, 2 × 95% EtOH, 1 × 80% EtOH, 1 × 70% EtOH, 3 × ddH2O; 3 min each. Heat induced epitope retrieval was then performed at 97°C for 10 min using the pH-9 Dako Target Retrieval Solution (Agilent, S236784-2) in a Lab Vision PT Module (Thermo Fisher Scientific). Slides were cooled to 65°C in the PT Module and then removed for equilibration to room temperature. A hydrophobic barrier was drawn around the tissue using the ImmEdge Hydrophobic Barrier pen (Vector Labs, 310018). Afterwards, endogenous peroxidase was inactivated using RNAscope Hydrogen Peroxide from the ACDBio RNAscope Multiplex Fluorescent Reagent Kit V2 (Biotechne, 323110), for 15 min at 40°C, followed by 2 × 2 min ddH2O washes. Coverslips were incubated overnight at 40°C (16 hrs) with RNAScope probes targeting human *C1QA* mRNA (Biotechne, 485451). Branch amplification was performed with Multiplex Amp 1, 2, 3 and HRP-C1 in the V2 kit: Amp1 30 min at 40°C, Amp2 15 min at 40°C, Amp3 30 min at 40°C, HRP-C1 15 min at 40°C, with 2 × 2 min 0.5 × RNAscope wash Buffer (Biotechne, 310091) washes between each steps. Coverslips were then incubated with TSA-Cy3 (Akoya Biosciences, NEL744001KT) in 1 × RNAscope TSA Buffer at a 1:50 dilution, for 15 min at room temperature in the dark, followed by 2 × 2 min 0.5 × RNAscope wash Buffer washing. The coverslips were then washed 2 × 5 min with 1× TBS-T, then subsequently blocked in Antibody Blocking Buffer (1 × TBS-T, 5% Donkey Serum, 0.1% Triton X-100, 0.05% Sodium Azide) for 1 hour. Antibody staining was next performed at 4°C overnight (16 hrs), with anti-CD15 (1:100 dilution, clone: MC480, Biolegend, 125602) and anti-CD68 (1:100 dilution, clone: D4B9C, Cell Signaling Technology, 76437T) in Antibody Dilution Buffer (1 × TBS-T, 3% Donkey Serum, 0.05% Sodium Azide). After staining, coverslips were washed 3 × 10 min with 1 × TBS-T, then incubated with secondary antibodies: AntiMouse-Cy7 (1:250, Biolegend, 405315) and Anti-Rabbit-Alexa647 (1:250, Thermo Fisher Scientific, A-21245) in Antibody Dilution Buffer for 30 min at room temperature. Coverslips were then washed 3 × 10 min with 1 × TBS-T, stained with Hoechst 33342 (1:10000 in 1 × TBS-T, Thermo Fisher Scientific, H3570) for 10 min at room temperature, and mounted with ProLong™ Diamond Antifade Mountant (Thermo Fisher Scientific, P36961).

Images were collected using a Keyence BZ-X710 inverted fluorescent microscope (Keyence, Inc) configured with 4 fluorescent filters (Hoechst, Cy3, Cy5 and Cy7), and a CFI Plan Apo l 20x/0.75 objective (Nikon). The Imaging setting was: 3 × 5 tile per tissue core, 5 Z-stacks acquired each FOV (best focused plane used), with High Resolution setting. The exposures were: 1/50s (Hoechst), 1/250s (Cy3), 1/8s (Cy5), and 6s (Cy7). Segmentation was performed with a local implementation of Mesmer (87), with weights downloaded from: https://deepcell-data.s3-us-west-1.amazonaws.com/model-weights/Multiplex_Segmentation_20200908_2_head.h5. Inputs of segmentation were Hoechst (nuclear) and *C1QA* + CD68 + CD15 (membrane). Signals from the images were capped at the 99.7th percentile, with prediction parameter `model_mpp` = 0.8. Features from single cells in segmented Keyence images were extracted based on the segmentation generated above, scaled by cell size, and written out as FCS files. Cells were filtered out if too large (CellSize > 500 pixels), too small (CellSize < 45 pixels) or limited in nuclear signal (Hoechst < 3500). The signal threshold of CD15, CD68 and *C1QA* positive cells were selected for each individual tissue core, and visually assessed to minimize false negative and false positive cells. Cells positive for CD68 and *C1QA* were annotated as *C1Q* High macrophages. The correlation of *C1Q* High macrophages between PANINI and CODEX experiments were calculated with the R function `cor()` with method `spearman`.

For spatial correlation analysis of C1QA expression in macrophages, the tissue core was divided into 100 subregions (a 10 × 10 grid), and the number of cells or *C1QA* signal level were summed in each individual region and plotted. Correlation was calculated with function `cor()` with method `spearman`.

#### Preprocessing and analysis of human PBMC datasets

CyTOF data measuring 33 proteins of PBMC from healthy human donors in Hartmann et al (94) was downloaded from flow-repository (‘FR-FCM-Z249, HD06_run1’). Cells were downsampled to 50,000, clustered using `Seurat` and manually annotated, and then a total of 38,866 annotated cells were used. CITE-seq data measuring 29 proteins of health human PBMC was retrieved from 10x genomics https://support.10xgenomics.com/single-cell-gene-expression/datasets/3.0.2/5k_pbmc_protein_v3?. Counts were normalized via CLR normalization with `Seurat` function `Normalizedata()`, then cells were clustered based on their protein features in `Seurat`. A total of 5,241 cells were annotated and used for matching. During matching, CITE-seq cells were used to match against CyTOF cells, where the input of CITE-seq cells were raw counts and the input of CyTOF cells were arcsine transformed with cofactor = 5. The MARIO parameters used were: `n_components_ovlp` = 10, `n_components_all` = 15, `sparsity` = 1000, `bad_prop` = 0.2, and `n_batch` = 1. Analysis was performed the same as previously described.

#### Preprocessing and analysis of cross species H1N1/IL-4 challenged datasets

Human H1N1 virus challenged data is the same as described in the previous section and the same set of cells were used as input to MARIO matching.

IL-4 stimulation cross-species CyTOF data is the same cross-species dataset as described in the previous section, using the same human or animal donors as described above (human: “7826”, “7718”, “2810”; Rhesus macaque: “D00522”, “D06022”, “D06122”; Cynomolgus monkey: “D07282”, “D07292”, “D07322”), and the whole blood cells stimulated with IL-4. Cells gated as Erythrocytes, Platelets and CD4+CD8+ cells from the paper (37) were excluded from downstream matching and analysis. Each individual dataset was randomly downsampled to 120,000 cells, arcsine transformed with cofactor = 5, and subsequently clustered with `Seurat` using all the markers, followed by manual annotation and removal of cells with ambiguous annotations. Total cell numbers for matching were 108,538 (human); 110,328 (rhesus macaque); 90,302 (cynomolgus monkey). During matching, human H1N1 challenged cells were matched against human, rhesus macaque and cynomolgus monkey IL-4 cells separately, and cells that matched to all three other datasets were used for downstream analysis. The MARIO parameters used: `n_components_ovlp` = 20, `n_components_all` = 15, `sparsity` = 1000, `bad_prop` = 0.1, `n_batch` = 4. Analysis was performed the same as previously described.

### Datasets benchmarking metrics and other methods

#### Benchmarking on the matching quality

Three scenarios were tested during the benchmarking process:

1. Sequentially dropping shared features between datasets, in order to test the robustness of the algorithm regardless of the antibody panel design.
2. Stimulated poor quality data by adding increasing levels of random noise to both datasets, in order to test the robustness of the algorithm in terms of over-integration. Gaussian random noise with mean 0 and standard deviation of 0.1, 0.3, 0.5, 0.7, 0.9, 1.1, 1.3, 1.5 was added to the normalized values of all protein channels.
3. Intentionally dropping cell types in the dataset being matched against, in order to test the robustness of the algorithm regardless of the cell type composition difference between datasets.

In all three scenarios described above, all other compared methods used the exact same set of cells tested by MARIO. For cross species data (related to Figure 3 and Figure S6) only H1N1 challenged human and X-species cynomolgus monkey were benchmarked.

The following metrics were used in the benchmarking process:

- *Matching accuracy*. Matching accuracy was calculated by the percentage of cells in *X* that have paired correctly with the same cell type in *Y*, based on the individual dataset’s cell type annotations.
- *Matching proportion*. Matching proportion was calculated by the percentage of cells in *X* that has a match in *Y* after quality control steps.
- *Structure alignment score*. Structure alignment score measures how much structural information is preserved after data integration. Let *D*_full_ be the matrix whose (*i, j*)-th entry is the Euclidean distance between the *i*-th row and the *j*-th row of *X*. Similarly, let *D*_partial_ be the matrix whose (*i, j*)-th entry is the Euclidean distance between the *i*-th row and the *j*-th row of the embedding of *X*. The structure alignment score for the *i*-th cell in *X* is defined as the Pearson correlation between the *i*-th row of *D*_full_ and the *i*-th row of *D*_partial_. The structure alignment score for *X* is then defined as the average of the scores over all cells in *X*. The structure alignment score for *Y* can be similarly obtained. The final structure alignment score is the average of the scores for *X* and *Y*.
- *Silhouette F1 score*. Silhouette F1 score has been described (31948481) and is an integrated measure of the quality of dataset mixing and information preservation. In brief, two preliminary scores `slt_mix` and `slt_clust` were obtained, and the Silhouette F1 score was calculated as 2· slt_mix · slt_clust*/*(slt_mix + slt_clust). Here, `slt_mix` is a measure of dataset mixing and is defined as one minus normalized Silhouette width with the label being dataset index, this is a measure of mixing; `slt_clust` is a measure of information preservation and is defined as the normalized Silhouette width with label being cell type annotations. All Silhouette widths were computed using the `silhouette()` function from R package `cluster`.
- *Adjusted Rand Index (ARI) F1 score*. ARI F1 score is an integrated measure of the quality of dataset mixing and information preservation (95). The definition is similar to that of Silhouette F1 score, except that we compute Adjusted Rand Index instead of the Silhouette width. All ARI scores were computed using the function `adjustedRandIndex()` in R package `mclust`.
- *Average mixing score*. Average mixing score is a measure of dataset mixing based on the Kolmogorov–Smirnov (KS) statistic. For each cluster, the subsets of cells corresponding to that cluster were extracted from the embeddings of *X* and *Y*, respectively. For each coordinate of the embeddings, one minus the KS statistic was computed. The mixing score for that cluster was then computed by taking the median of one minus the KS statistic for each coordinate. The average mixing score is defined as the average of mixing scores over all clusters.
- *Error avoidance score*. Error avoidance score measures the performance of the quality control process and is specific to the benchmarking scenario 3 (intentionally dropping cell types). For each cell type dropped, the corresponding error avoidance score is defined as 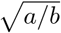, where *a* is the number of cells in *X* that are of that type and have survived the quality control process (i.e., a match involving that cell type has occurred), and *b* is the total number of cells of that type *X*. Higher value of this score indicates that erroneous matching towards deleted cells types has been avoided more.

During benchmarking, all datasets were downsampled. The Bone marrow dataset (Figure 2) was downsampled to 40,000 cells (8000 and 32,000 for *X* and *Y*); the PBMC dataset (Figure S3) was downsampled to 25,000 cells (5000 and 20,000 for *X* and *Y*); the X-Species H1N1/IFN-gamma dataset (Figure 3) was downsampled to 40,000 cells (8000 and 32,000 for *X* and *Y*); the X-Species H1N1/IL-4 dataset (Figure S6) was downsampled to 40,000 cells (8,000 and 32,000 for *X* and *Y*); and the Murine spleen dataset (Figure 4) downsampled to 25,000 cells (5000 and 20,000 for *X* and *Y*). All methods used the same set of cells.

Parameters used for benchmarking are as follows. For benchmarking of MARIO, we used a consistent set of parameters across all datasets: `n_components_ovlp` = 10 (or the maximum number available); `n_components_all` = 20 (or the maximum available), `sparsity` = 5000, `bad_prop` = 0.1, `n_batch` = 1. For other methods, the input of data were all values normalized per feature within each dataset (except Liger where their own custom normalization is required). Only mNN-based methods (Scanorma, Seurat, fastMNN) were included in the comparison of matching accuracy and matching proportion. All methods used default parameters, using available shared features. For computation of SAM, ASW, ARI and avgMix, the first 20 (or maximum available) components of MARIO CCA scores or reduced values from other methods were used. For visualization, t-SNE plots were produced using the first 10 components for all methods.

#### Benchmarking on time and memory usage

Time and memory usage of MARIO on the datasets presented in Figure 2, 3, 4 were evaluated. The full pipeline MARIO time usage (including initial and refined matching; best interpolation finding; joint regularized filtering; CCA calculation) was measured with the default parameters, with increasing amount of cells (50,000 cell max), and ratio of *X* and *Y* set to 1:4 (e.g. at total of 20,000 cells, *X* has 4000 cells and *Y* has 16,000 `memory_profiler`. The influence of sparsity level used cells). The MARIO matching time usage (only including intial and refined matching) was measured with the same settings, but with three different sparsity levels: (1) minimal sparsity calculated by MARIO; (2) maximal sparsity (i.e., fully dense matching without sparsification); (3) “medium” sparsity which is in the middle point between minimal and maximum. The MARIO memory usage was measured with the same settings as the time evaluation, but the maximum number was set to 100,000 cells. The peak memory usage was measured by the function profile in the python package on MARIO matching accuracy was evaluated by inputting different levels (between minimal and maximal sparsity detected by MARIO). A total of 50,000 cells were used for each dataset with a ratio between *X* and *Y* being 1:4.

## ACKNOWLEDGEMENTS

We thank Sean Bendall, Scott Rodig and members of the Nolan and Jiang labs for helpful discussions. B.Z. is supported by a Stanford Graduate Fellowship. This work was funded in part by grants from the National Institutes of Health R01AI149672 (S.J., G.P.N.), the Bill & Melinda Gates Foundation INV-002704 (S.J., G.P.N.), OPP1113682 (G.P.N.), COVID-19 Pilot Award (S.J., D.R.M., G.P.N.), the Fast Grant Funding for COVID-19 Science (S.J., D.R.M., G.P.N.), the Botnar Research Centre for Child Health Emergency Response to COVID-19 grant (S.J., D.R.M., G.P.N., M.S.M., and A.T.), the US Food and Drug Administration Medical Countermeasures Initiative contracts HHSF223201610018C and 75F40120C00176 (G.P.N.), the Parker Institute for Cancer Immunotherapy (G.P.N.), and the Rachford and Carlota A. Harris Endowed Professorship (G.P.N.). This article reflects the views of the authors and should not be construed as representing the views or policies of the FDA, NIH, BMGF, Botnar Foundation or other institutions who provided funding.

## AUTHOR CONTRIBUTIONS

Conceptualization: B.Z., S.C., Z.M., G.P.N., S.J.

Algorithm Development and Implementation: S.C., B.Z., Z.M.

Analysis: B.Z., S.C., Y.B., H.C., I.T.L., Y.G., S.J.

Contribution of Key Reagents and Tools: N.M., G.V., D.R.M., A.T., M.M.

Supervision: S.J., G.P.N., Z.M.

Both B.Z. and S.C. contributed equally and have the right to list their name first in their CV.

## Supplementary Figures

**Figure S1:**
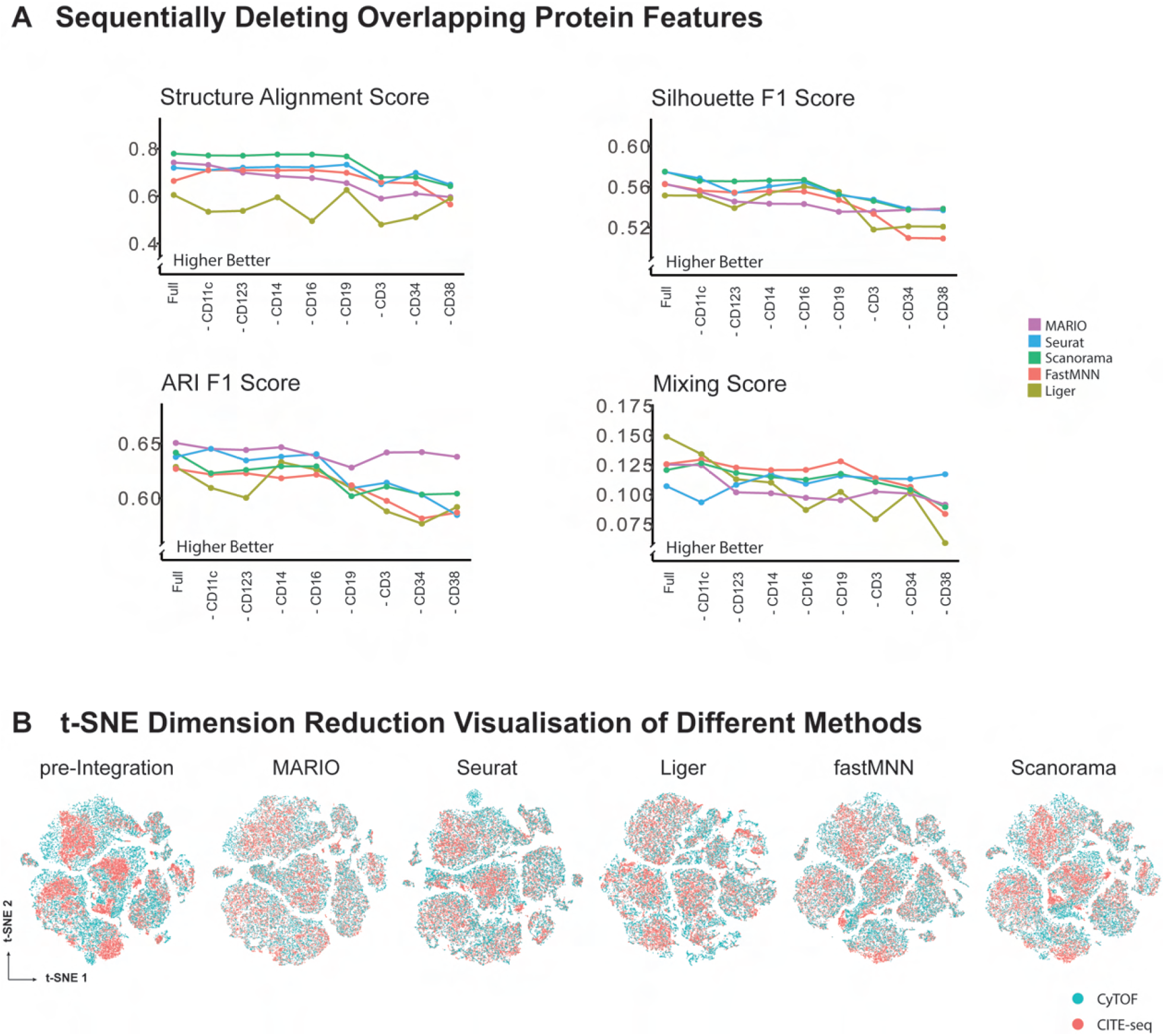
Performance of matching and integration on bone marrow cells in relationship to Figure 2. Comparison of MARIO and other mNN methods, related to Figure 2. **(A)** Performance of matching and integration during sequentially dropping of shared protein features. The tested parameters shown here are: average Structure alignment score, Silhouette F1 score, Adjusted Rand Index F1 score and average Mixing score. **(B)** t-SNE plots visualizing pre-integation and post-integration results with different methods. For methods other than MARIO, only shared features were used during integration.

**Figure S2:**
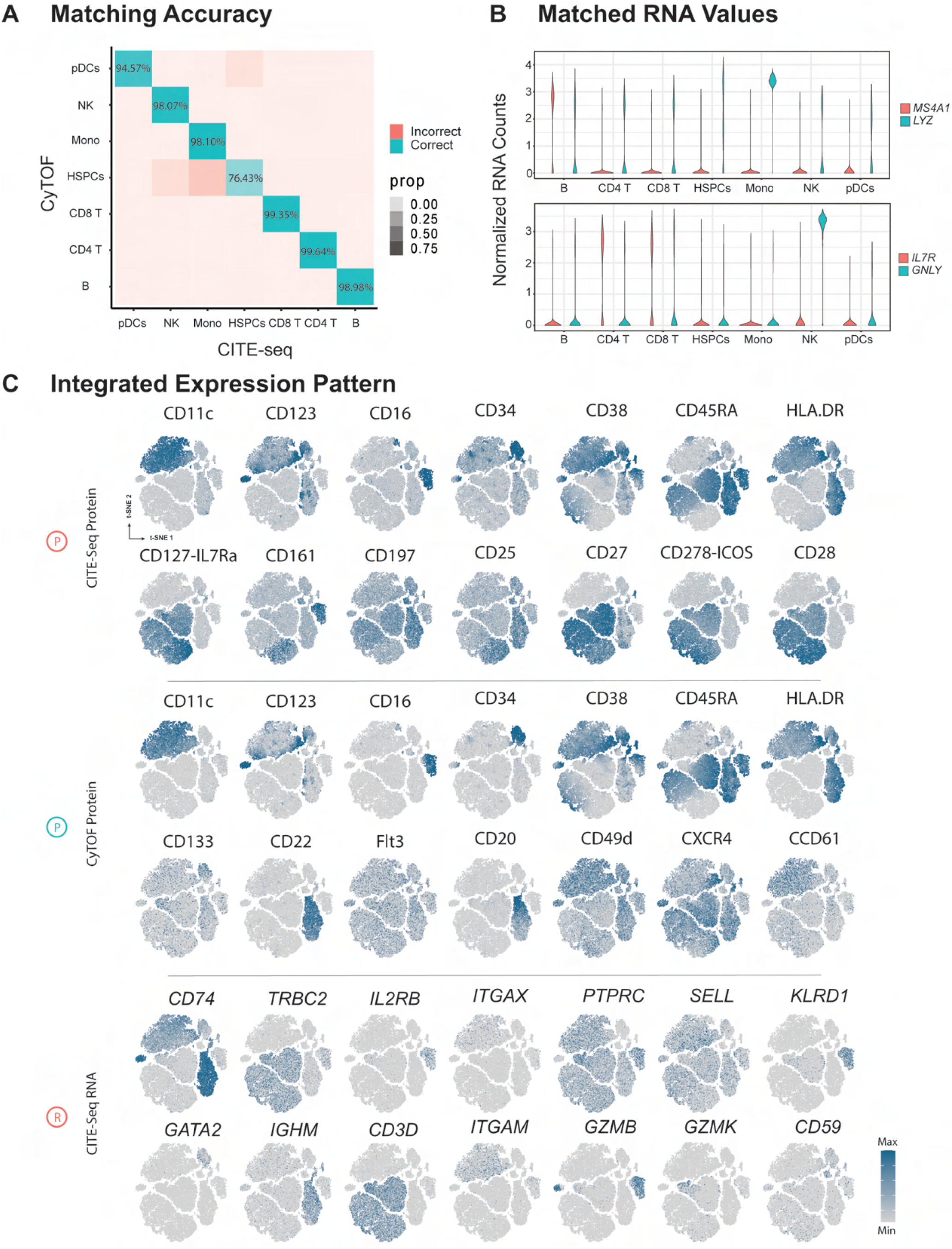
Matching and integration of cross-modality CyTOF and CITE-seq bone marrow data with MARIO, related to Figure 2. **(A)** Confusion matrix with MARIO cell-cell matching accuracy (balanced accuracy) across cell types. **(B)** Violin plots of normalized RNA counts among different MARIO matched CITE-seq and CyTOF cell types. **(C)** t-SNE plots of the matched cells with protein/RNA expression levels overlaid as an extension of Figure 2G.

**Figure S3:**
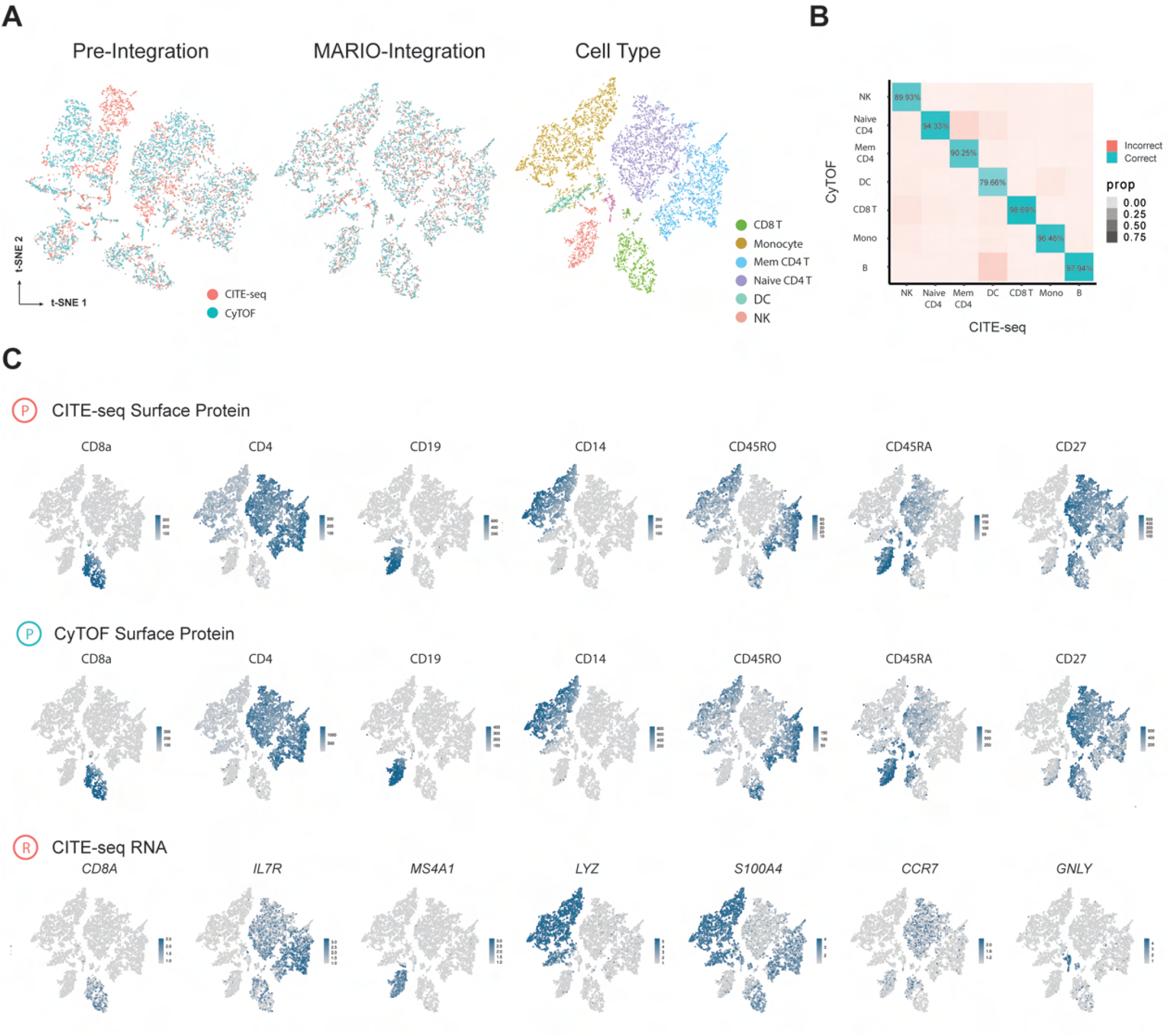
Matching and integration of cross-modality CyTOF and CITE-seq PBMC data with MARIO. MARIO integration of human PBMCs as measured by CyTOF and CITE-seq. **(A)** t-SNE plots of the PBMC CITE-seq and CODEX cells, pre-integration (left) and MARIO integrated (middle and right), colored by dataset of origin (left and middle) or colored by cell types (right). **(B)** Confusion matrix with MARIO cell-cell matching accuracy (balanced accuracy) across cell types. **(C)** t-SNE plots of the matched cells with protein or RNA expression levels overlaid.

**Figure S4:**
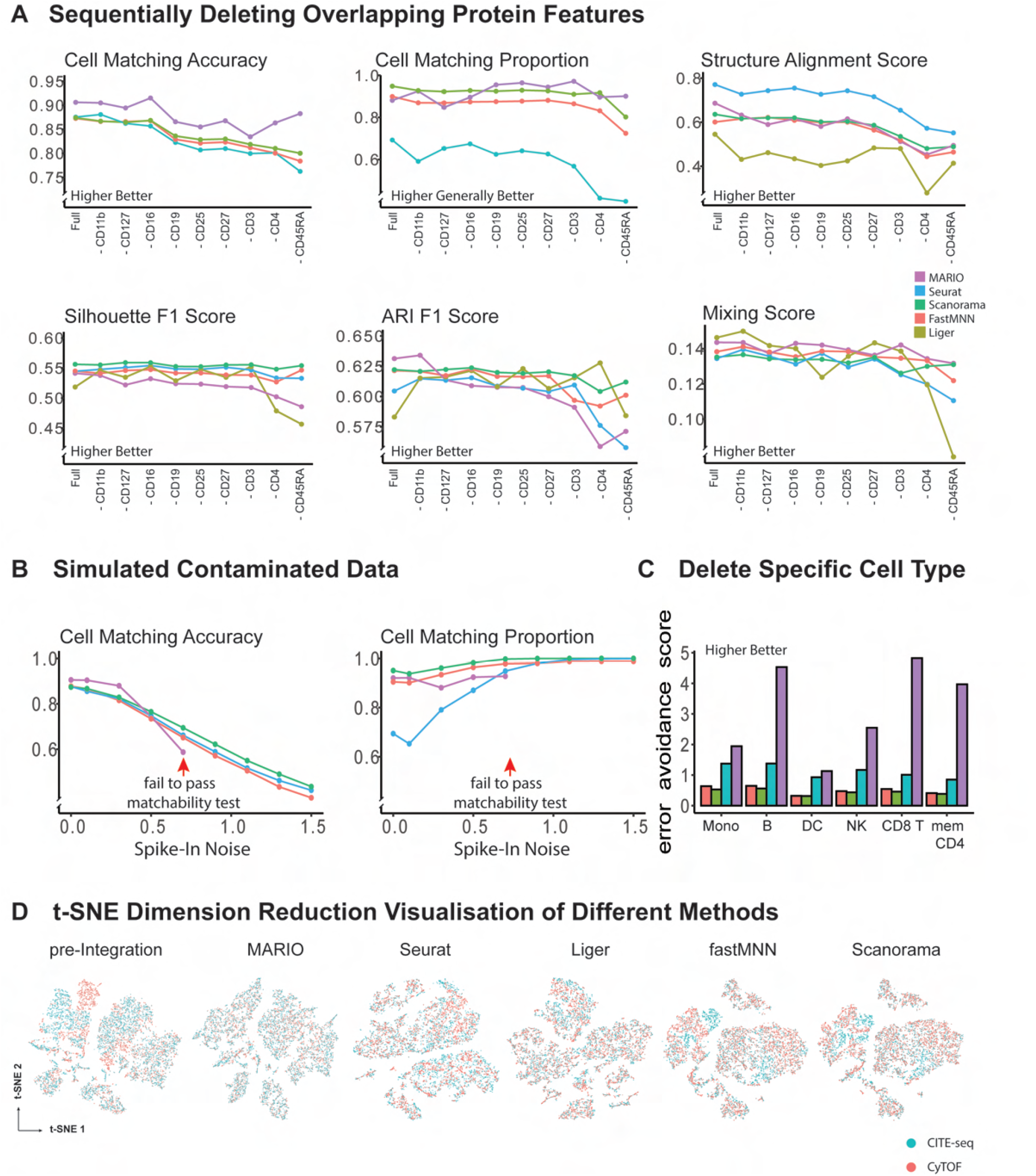
Performance of matching and integration on PBMCs in relationship to Figure S3. **(A)** Performance of matching and integration during sequentially dropping of shared protein features. The tested parameters are: cell-cell matching accuracy, proportion of cell in *X* matched, average Structure alignment score, Silhouette F1 score, Adjusted Rand Index F1 score and average Mixing score. **(B)** Testing algorithm stringency between different methods. Increasing amounts of random spike-in noise was added to the data, and the matching accuracy and proportion of cells matched to *X* were quantified. MARIO matchability test automatically suspended forced matching of inappropriate data due to poor quality here. **(C)** Testing algorithm stringency among different methods. Single-cell types in *Y* were deleted before matching to *X*. The proportion of cells belonging to the deleted cell type in matched *X* cells were used to calculate the erroneous avoidance score. **(D)** t-SNE plots visualizing pre-integation and post-integration results with different methods.

**Figure S5:**
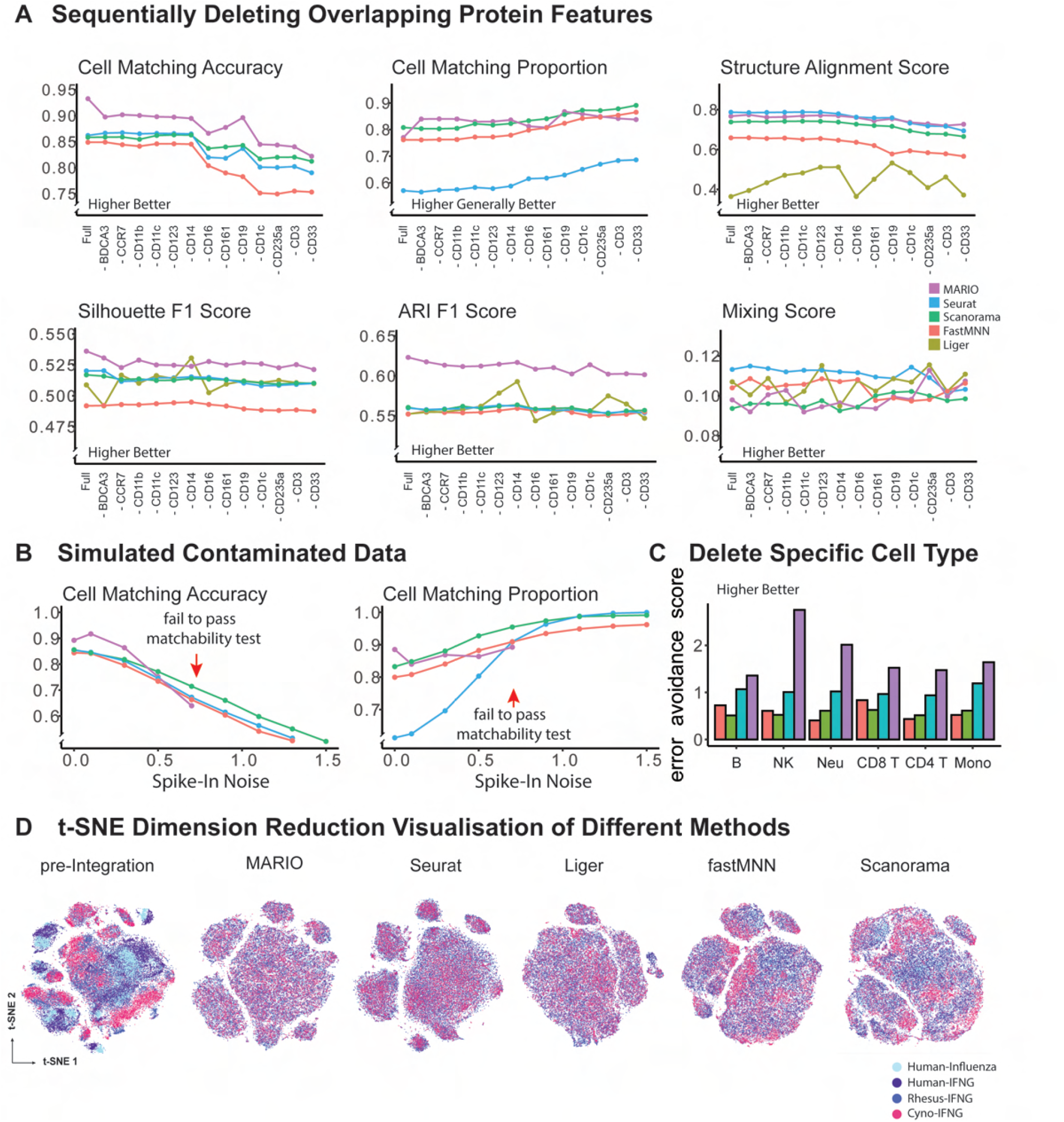
Performance of matching and integration on cross-species whole blood cells CyTOF data in Figure 3. **(A)** Performance of matching and integration during sequentially dropping of shared protein features. The tested parameters are: cell-cell matching accuracy, proportion of cell in *X* matched, average Structure alignment score, Silhouette F1 score, Adjusted Rand Index F1 score and average Mixing score. **(B)** Testing algorithm stringency between different methods. Increasing amounts of random spike-in noise was added to the data, and the matching accuracy and proportion of cells matched to *X* were quantified. MARIO matchability test automatically suspended forced matching of inappropriate data due to poor quality here. **(C)** Testing algorithm stringency among different methods. Single-cell types in *Y* were deleted before matching to *X*. The proportion of cells belonging to the deleted cell type in matched *X* cells were used to calculate the erroneous avoidance score. **(D)** t-SNE plots visualizing pre-integation and post-integration results with different methods.

**Figure S6:**
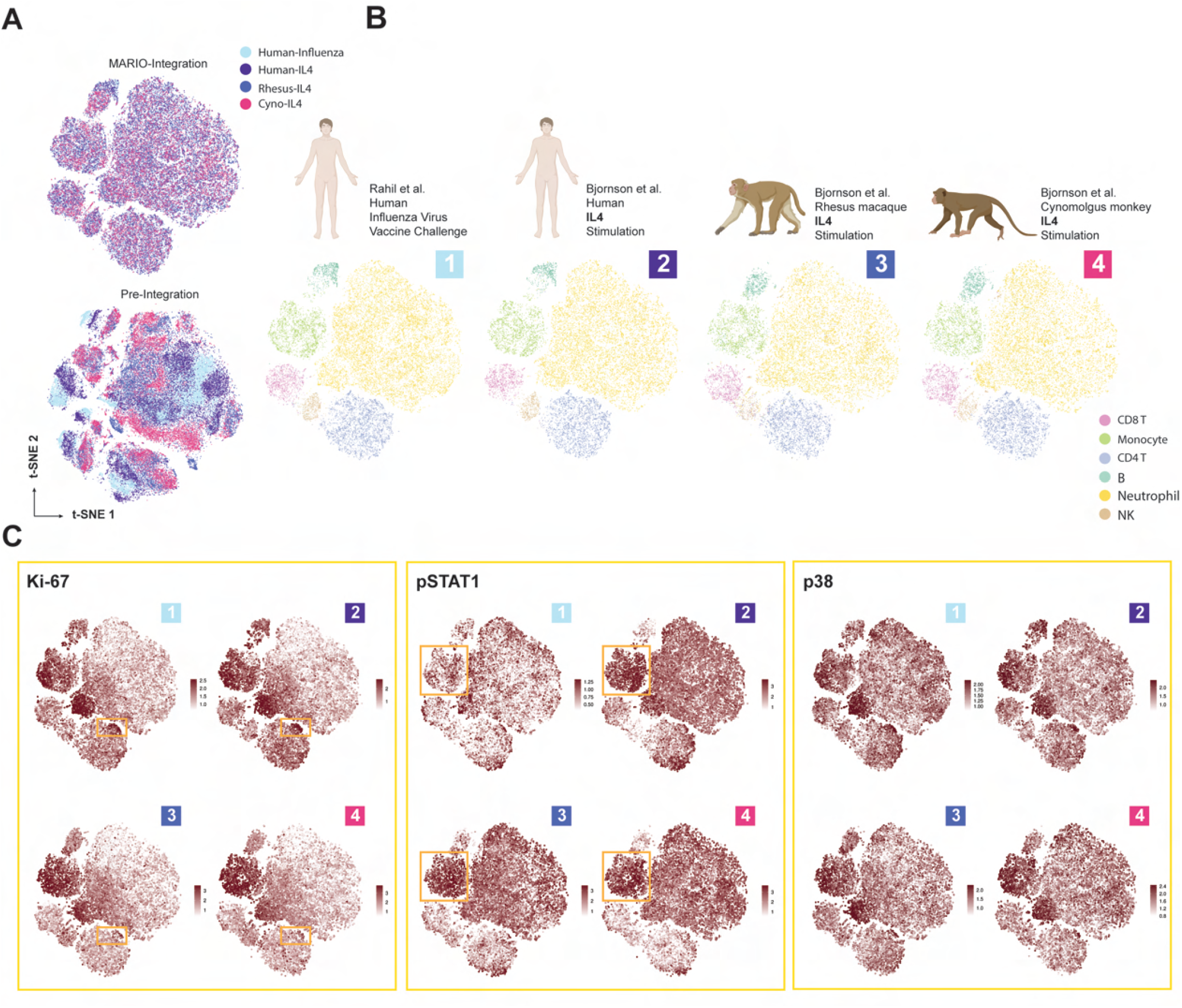
Cross-species H1N1 Challenge and IL-4 integrative analysis with MARIO. MARIO integration of human, rhesus macaque and cynomolgus monkey whole blood cells from a H1N1 challenge study or IL-4 stimulation. **(A)** t-SNE plots of the four datasets, pre-integration and post MARIO-integration as colored by dataset of origin. **(B)** t-SNE plots of each individual dataset, colored by cell type annotation. **(C)** t-SNE plots with expression levels of Ki-67, STAT1 and p38 across four datasets.

**Figure S7:**
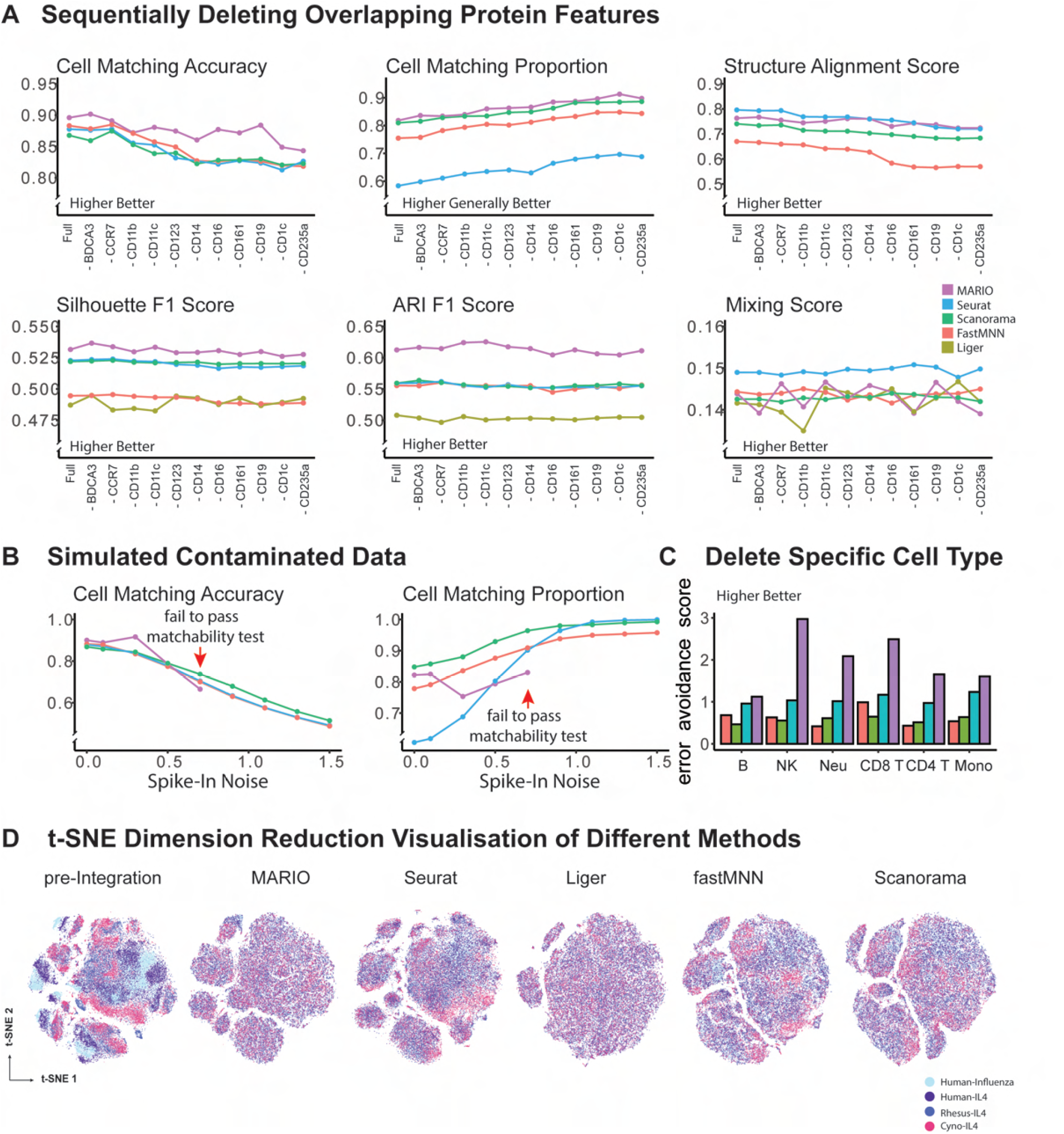
Performance of matching and integration on cross-species whole blood cells CyTOF data in Figure S6. **(A)** Performance of matching and integration during sequentially dropping of shared protein features. The tested parameters are: cell-cell matching accuracy, proportion of cell in *X* matched, average Structure alignment score, Silhouette F1 score, Adjusted Rand Index F1 score and average Mixing score. **(B)** Testing algorithm stringency between different methods. Increasing amounts of random spike-in noise was added to the data, and the matching accuracy and proportion of cells matched to *X* were quantified. MARIO matchability test automatically suspended forced matching of inappropriate data due to poor quality here. **(C)** Testing algorithm stringency among different methods. Single-cell types in *Y* were deleted before matching to *X*. The proportion of cells belonging to the deleted cell type in matched *X* cells were used to calculate the erroneous avoidance score. **(D)** t-SNE plots visualizing pre-integation and post-integration results with different methods.

**Figure S8:**
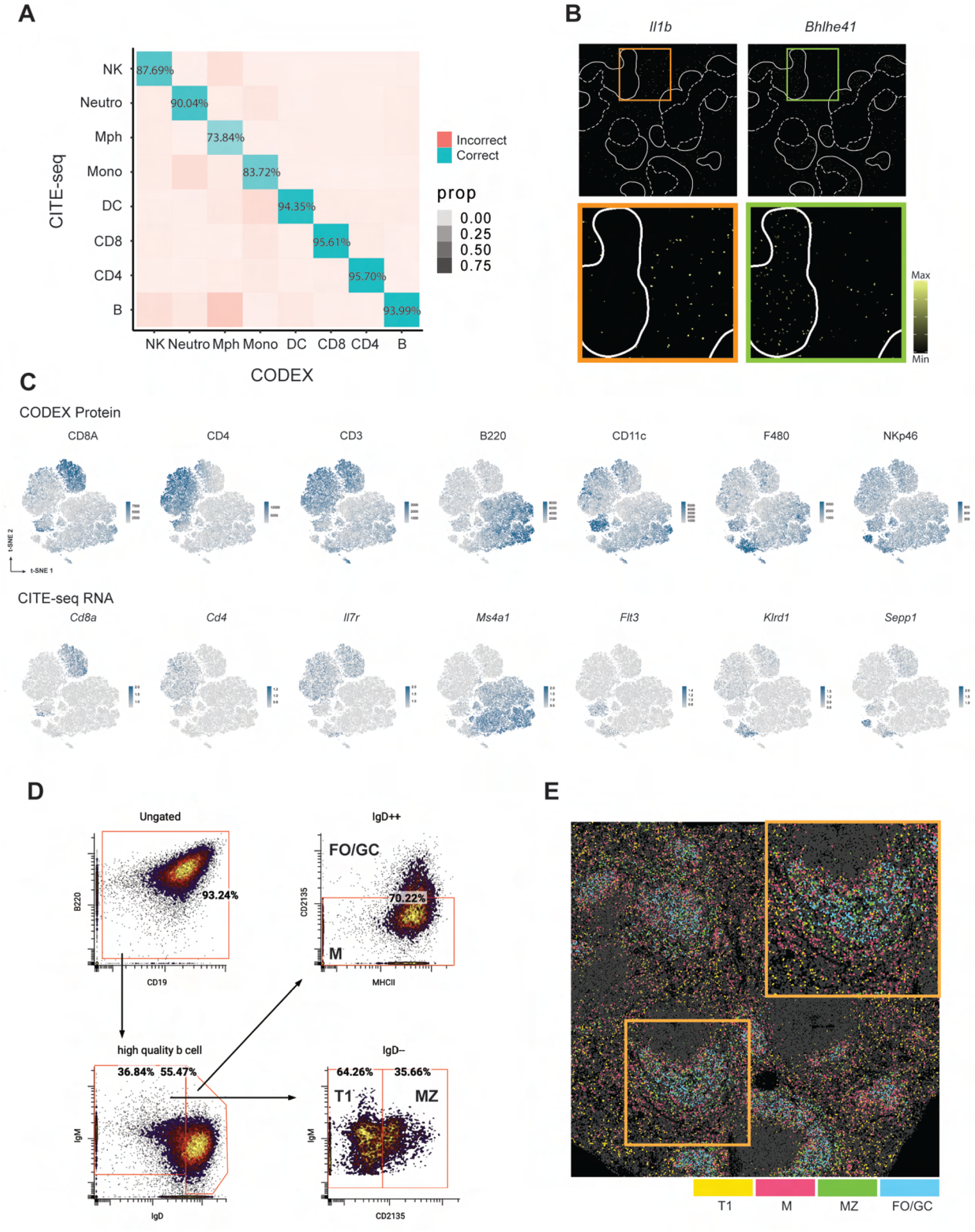
MARIO integrative analysis of CODEX and CITE-seq for spatial multi-omics. Related to Figure 4. **(A)** Confusion matrix with MARIO cell-cell matching accuracy (balanced accuracy) across cell types for matched CITE-seq or CODEX cells. **(B)** A pseudo-colored murine spleen section showing the localization of transcripts (*Il1b* and *Bhlhe41*) inferred from CITE-seq. The white outline demarcates the white pulp. **(C)** t-SNE plots of MARIO integrated murine spleen CITE-seq and CODEX cells, overlaid with matched CODEX protein and CITE-seq RNA expression levels. **(D)** Gating strategy of CODEX B cell subtypes (T1, MZ, M, FO/GC B cells) using CODEX single-cell protein expression. **(E)** A pseudo-colored murine spleen section colored by the annotation of CODEX B cell subpopulations, gated as previously described in (D).

**Figure S9:**
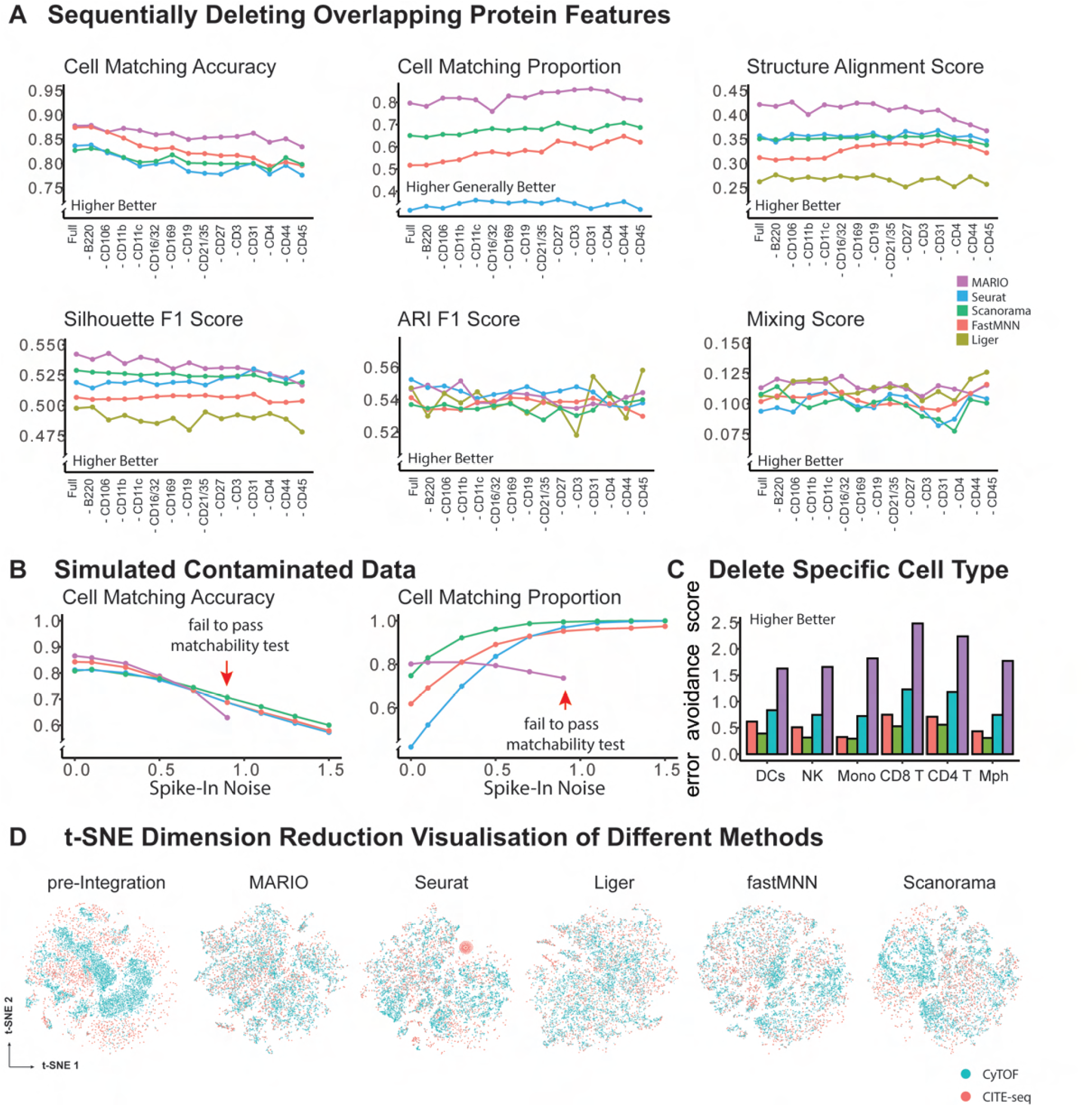
Performance of matching and integration on murine spleen cells in Figure 4. **(A)** Performance of matching and integration during sequentially dropping of shared protein features. The tested parameters are: cell-cell matching accuracy, proportion of cell in *X* matched, average Structure alignment score, Silhouette F1 score, Adjusted Rand Index F1 score and average Mixing score. **(B)** Testing algorithm stringency between different methods. Increasing amounts of random spike-in noise was added to the data, and the matching accuracy and proportion of cells matched to *X* were quantified. MARIO matchability test automatically suspended forced matching of inappropriate data due to poor quality here. **(C)** Testing algorithm stringency among different methods. Single-cell types in *Y* were deleted before matching to *X*. The proportion of cells belonging to the deleted cell type in matched *X* cells were used to calculate the erroneous avoidance score. **(D)** t-SNE plots visualizing pre-integation and post-integration results with different methods.

**Figure S10:**
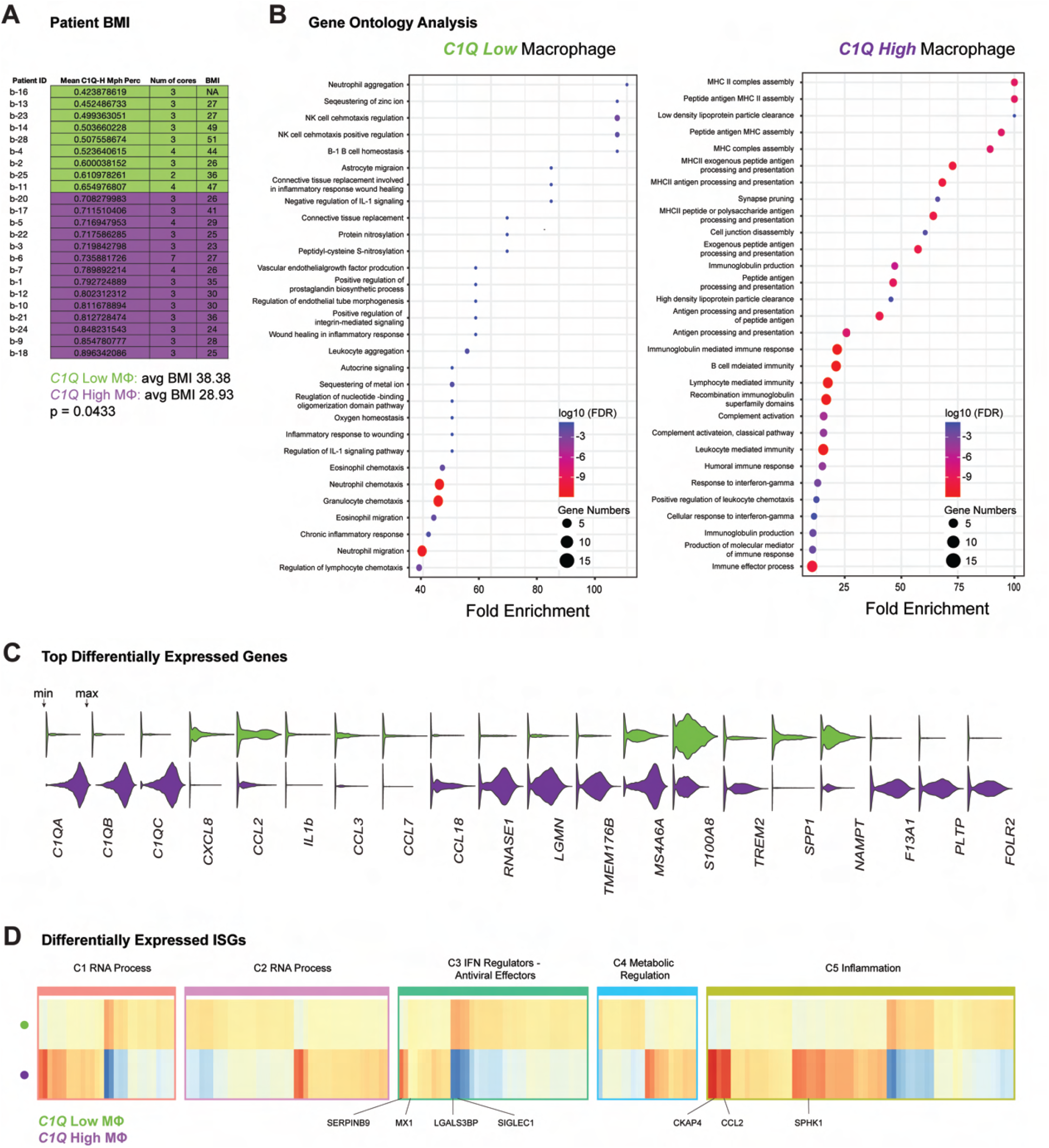
MARIO analysis on COVID-19 lung tissue and BALF cells. Related to Figure 5, part 1. **(A)** A table showing MARIO predicted *C1Q* high macrophages as a percentage of total macrophages in each patient, and their BMI values. P-values calculated using the student’s t-test. **(B)** GO term analysis for transcriptional programs enriched in *C1QA* low (left) and *C1QA* high macrophages (right). **(C)** Violin plots of selected genes from the top 50 differentially expressed genes (p-adjust < 0.05) for *C1Q* low (green) or *C1Q* high (magenta) macrophages. **(D)** A heatmap representation of differentially expressed ISGs among C1QA low (up) or C1QA low macrophages (down). Genes are categorized into 5 previously described classes of biological pathways (see Materials and Methods).

**Figure S10:**
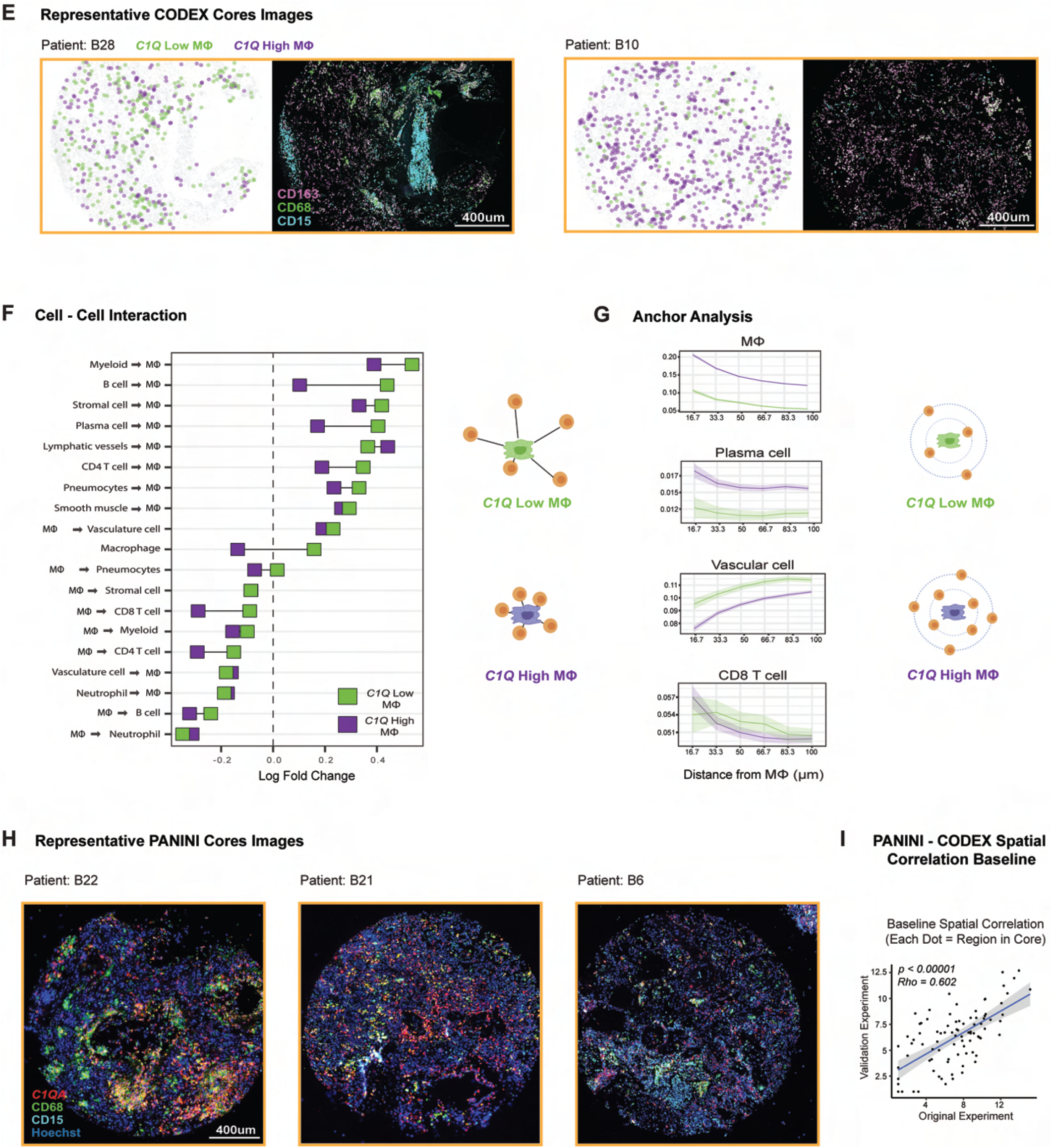
MARIO analysis on COVID-19 lung tissue and BALF cells. Related to Figure 5, part 2. (**E**) Additional representative CODEX images of COVID-19 lung tissue cores for patients with **C1Q** low (green) and high (magenta) macrophage locations. CD163, CD68 and CD15 antibody staining are shown on the right of each image. (**F**) The pairwise cell distances between C1Q high low (green) or (magenta) macrophages to other cell types, as an enrichment over the permutated background distribution. Only interactions that passed a statistical test (p<0.05) for both macrophage subgroups conditions are shown. Squares that are toward the left indicate interactions that are closer than expected, and those toward the right indicate interactions that are further apart than expected. (**G**) Anchor plots of average cell type fractions around C1Q low (green) or **C1Q** high (magenta) macrophages. The thick colored lines represent the means, and lighter regions around these lines depict the 95% confidence interval. The macrophages are anchored at 0 μm, and the plot ends at a 100 μm radial distance from the anchored macrophages. (**H**) Representative images of COVID-19 lung tissue cores in the PANINI validation experiment, stained with **C1QA**, CD68, CD15 and Hoechst. (**I**) Spatial correlation of cell density in each 10 × 10 region of the same tissue core between CODEX experiment and PANINI validation to determine the baseline correlation between the tissue sections for CODEX and PANINI (P-value and Correlation calculated via Spearman Ranked Test).

**Figure S11:**
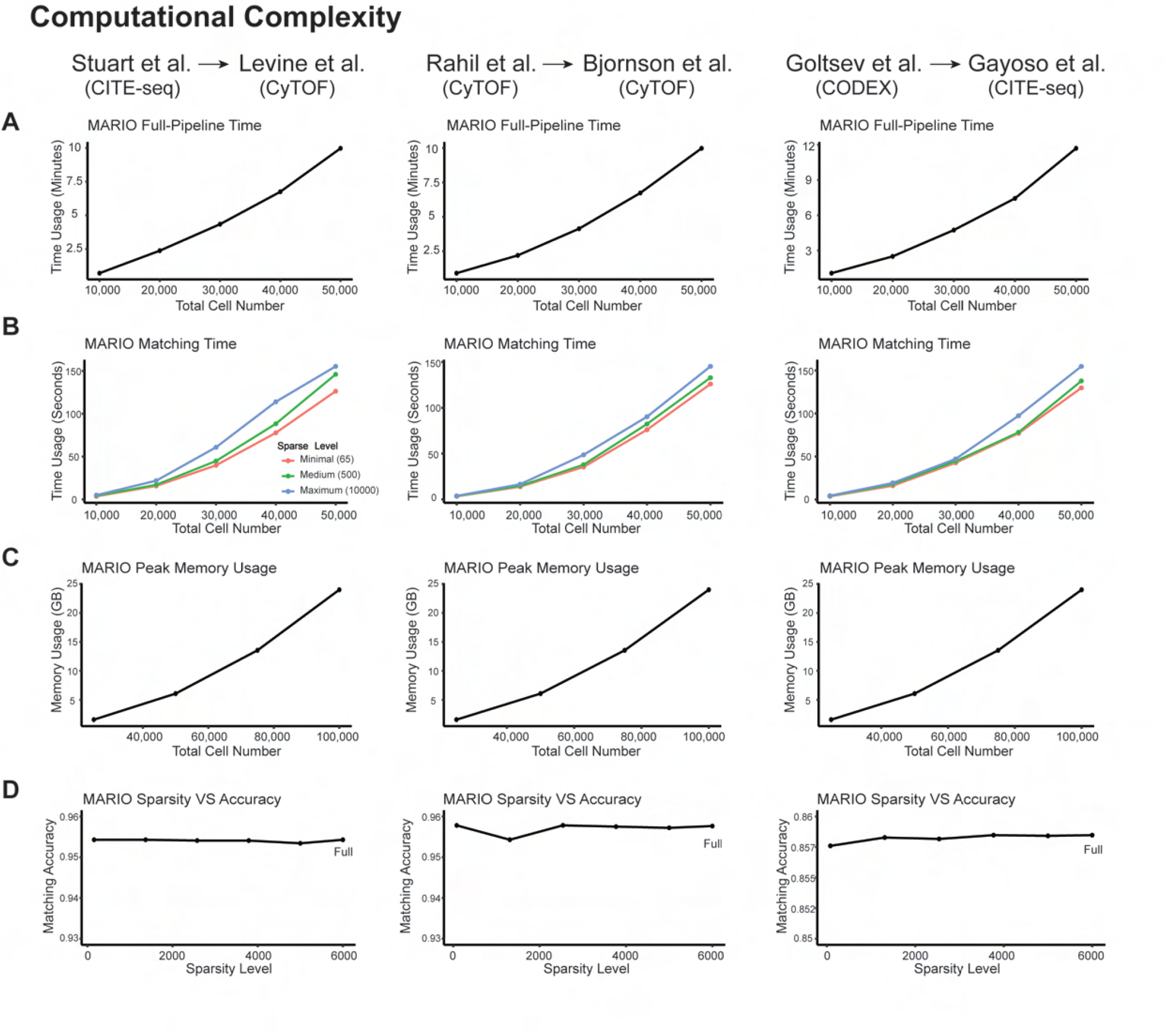
Figure S11 Computational complexity. **(A)** Run time for full MARIO pipeline (Initial and refined matching; Finding the best interpolation; Joint regularized filtering; CCA calculation) across different datasets. **(B)** Run time for MARIO matching steps (total time for initial and refined matchings) across different datasets. The ratio of *X* and *Y* was set as 1:4 (eg. at a total of 20,000 cells, *X* has 4000 cells and *Y* has 16,000 cells). Three sparsity levels were shown in the figures, which are 1: ‘Minimal’ sparsity calculated by MARIO. 2: ‘Maximum’ sparsity, same as using dense data. 3: ‘Medium’ sparsity which is the level in the middle between minimal and maximum. **(C)** Peak memory usage when running the full MARIO pipeline across different datasets. The ratio of *X* and *Y* was set as 1:4. **(D)** Matching accuracy with different levels of sparsity for MARIO. Total of 50,000 cells were used, where the ratio of *X* and *Y* was set as 1:4.

